# Complex multiple introductions drive fall armyworm invasions into Asia and Australia

**DOI:** 10.1101/2022.06.11.495773

**Authors:** R Rane, TK Walsh, P Lenancker, A Gock, TH Dao, VL Nguyen, TN Khin, D Amalin, K Chittarath, M Faheem, S Annamalai, SS Thanarajoo, YA Trisyono, S Khay, J Kim, L Kuniata, K Powell, A Kalyebi, MH Otim, K Nam, E d’Alençon, KHJ Gordon, WT Tay

## Abstract

The fall armyworm (FAW) *Spodoptera frugiperda* is thought to have undergone a rapid ‘west-to-east’ spread since 2016 when it was first identified in western Africa. Between 2018 and 2020, it was also recorded from South Asia (SA), Southeast Asia (SEA), East Asia (EA), and Pacific/Australia (PA). Population genomic analyses enabled the understanding of pathways, population sources, and gene flow in this notorious agricultural pest species. Using neutral single nucleotide polymorphic (SNP) DNA markers, we detected genome introgression that suggested most populations were overwhelmingly C- and R-strain hybrids. SNP and mitochondrial DNA markers identified multiple introductions that were most parsimoniously explained by anthropogenic-assisted spread, i.e., associated with international trade of live/fresh plants and plant products, and involved ‘bridgehead populations’ in countries to enable successful pest establishment in neighbouring countries. Distinct population genomic signatures between Myanmar and China do not support the ‘African origin spread’ nor the ‘Myanmar source population to China’ hypotheses. Significant genetic differentiation between populations from different Australian states supported multiple pathways involving distinct SEA populations. Our study identified Asia as a biosecurity hotspot and a FAW genetic melting pot, and demonstrated the use of genome analysis to disentangle preventable human-assisted pest introductions from unpreventable natural pest spread.

## Introduction

Movements of exotic arthropods are increasingly being linked with global agricultural and horticultural trade (Bedford 1980, Lopes-da-Silva et al. 2014, Tonnang et al. 2015, Tay et al. 2017, Elfekih et al. 2018, Arnemann et al. 2019, Europhyte <u>2018</u>), tourism (e.g., Early et al. 2018), and as hitchhiker pests on aircraft (Porter and Hughes 1950) and vessels (e.g., Paini et al. 2018). These exotic arthropods can have significant negative impact on agricultural production (De Barro et al. 2011, Tay et al. 2013, Pozebon et al. 2020, Haile et al. 2021), and the environments (e.g., Bedford 1980, O’Dowd et al. 2003). Good phytosanitary practice together with robust biosecurity preparedness can lower the risk and help mitigate the effects of these invasive species. Invasive arthropods, especially species of agricultural importance, may also introduce novel genotypes (e.g., insecticide resistance genes and alleles; Anderson et al. 2018, Walsh et al. 2018, Tay and Gordon 2019) into related native populations through introgression and hybridisation, or into established populations of invasive species, thereby increasing their genomic diversity and further complicating pest management (e.g., Anderson et al. 2018, Valencia-Montoya et al. 2020).

The fall armyworm (FAW; *Spodoptera frugiperda*), a highly polyphagous noctuid moth native to the tropical and subtropical regions of the Americas, was officially reported in Western Africa in early 2016 (Goergen et al. 2016), followed by rapid detections across Sub-Saharan Africa (e.g., Nagoshi et al. 2018, Otim et al. 2018b, Botha et al. 2019) by February 2018. In Asia, its occurrence was first reported in South Asia (SA) in India in May 2018, and by December 2018, it had been reported in Bangladesh, Sri Lanka, and in Myanmar (i.e., Southeast Asia, SEA), and in China (i.e., East Asia, EA) by January 2019. In SEA, throughout 2019, detection of FAW was reported from Indonesia (Trisyono et al. 2019), Laos, Malaysia, Vietnam (Hang et al. 2020), the Philippines (Navasero et al. 2019), Cambodia (S Khay pers. ob.), and in EA from the Republic of Korea (Seo et al. 2019, Kim et al. 2021) and Japan (<u>FAO 2021</u>) in July 2019. The moth’s strong flight ability (e.g., Rose et al. 1975, Sparks 1979, Jones et al. 2019) and potential to contaminate certain agricultural trade commodities (e.g., Early et al. 2018) leads to its success as a pest and its detection in increasing number of countries.

In the native range, FAW is conventionally divided into two host races, the corn (C-strain) and the rice (R-strain) preferred individuals. This distinction is evidenced by phenotypic and genotypic differences between the two. This host separation is thought to be an example of incipient speciation and has been primarily seen in North America. However, even in the native range, particularly in the tropics, this distinction seems to break down. Several studies have shown that, at the genetic level, the invasive populations are hybrids (e.g., Zhang et al. 2020; Tay et al. 2021b, 2022d).

In early 2020, FAW was detected in Pacific/Australia (PA) in Papua New Guinea (Bourke and Sar 2020, Tay et al. 2022a) and in Australia, FAW was first detected in the Erub and Saibai Islands in the Torres Strait in January 2020, and by early February it was confirmed from maize fields in Queensland (QLD), followed by detection in Western Australia (WA), Northern Territory (NT), New South Wales (NSW), Victoria (VIC), and as of April 2021, it was also reported from Tasmania, as well as Norfolk Island in the Pacific, and most recently (March 2022) also in New Zealand. The FAW is now established across northern Australia and has been reported in a number of crops including sorghum and sugarcane. Since reported by Goergen et al. (2016), the rapid detections of FAW across the sub-Saharan African nations followed by Asian (i.e., SA, SEA, EA) detections led to the general acceptance in the scientific and agricultural sectors that the pest spread eastward from an initial incursion into Africa (Day et al. 2017). This perception fits the reported patterns of detection but does it really fit the actual movement of this species across the Old World?

Studies based on nuclear DNA markers involving African (e.g., Schlum et al. 2021, Tay et al. 2022d) and Asian (e.g., Jiang et al. 2022) invasive populations identified gene flow patterns and signatures that suggested the FAW’s recent spread was due to multiple introductions, with invasive populations likely to have moved from the east (i.e., EA/SEA) to Africa (e.g., Tay et al. 2022d). Genomic analyses of Chinese and African FAW populations by Gui et al. (2020) also detected signatures that were best explained by independent introduction events in EA, and a likely east-to-west movement of the FAW. The authors’ conclusion however, nevertheless maintained the prevailing axiom that FAW arrived in China from western African, with its most recent origin to the Yunnan Province being from the neighbouring country of Myanmar (Sun et al. 2019, Zhang et al. 2019, Gui et al. 2020, Li et al. 2020).

Understanding the spread patterns (e.g., single invasive bridgehead effect (Guillemaud et al. 2011) vs. mass dispersion (Wilson et al. 2009)) and frequencies of introduction (e.g., single introduction (Cock et al. 2017, Nagoshi et al. 2018, Nagoshi et al. 2019b, Gui et al. 2020) vs. multiple introductions (Schlum et al. 2021, Tay et al. 2021a, Tay et al. 2021b, Jiang et al. 2022, Tay et al. 2022a, Tay et al. 2022c)) of the current invasive FAW populations in Africa, Asia, and Pacific/Australia will have significant implications for the future management of this pest, especially with respect to the delimitation between human-assisted introductions (i.e., preventable via behavioural change) vs. climatic factor-assisted natural migration (i.e., difficult to prevent/not preventable), insecticide resistance management (Tay et al. 2021a, Tay et al. 2022c), and climate (i.e., hot and cold) adaptation and tolerance (e.g., Foster and Cherry 1987, Keosentse et al. 2021, Yang et al. 2021, Zhang et al. 2021), highlighting most likely agents of spread. It also highlights that evaluating possible future incursion routes makes it necessary to identify weak points in the phytosanitary risk assessments, to enable future targeted monitoring at the right places, and to critically assess official reporting and announcement dates. The implications are significant, such as in reverse simulation studies (e.g., Sun et al. 2019, Wu Q et al. 2019, Li et al. 2020, Qi et al. 2021) that relied on reported detections dates to identify potential wind patterns for explaining pest dispersal patterns, but have not incorporated appropriate genomic/genetic data to ensure consistency between multiple datasets, and could result in significant biosecurity concerns relating to anthropogenic-assisted introductions not being addressed due to the perceived difficulty of preventing pest spread via natural migration.

In this study, we aim to apply genome-wide single nucleotide polymorphic (SNP) loci to: (i) determine the genomic diversity of invasive populations of FAW and the likelihood of multiple introduction events, and (ii) determine the area of origin and introduction pathways of FAW populations in Australia. This information can then be used to delineate between human-assisted vs. climate-assisted migration patterns of FAW in SEA and PA regions and inform risk assessments for future incursions of both natural and anthropogenic spread.

## Results

### Strain assessment

We did not detect any C-strain individual following analysis of 138 fully assembled mitochondrial DNA genomes (mitogenomes) from Australian samples. Our results are similar to the finding by Piggott et al. (2021) who detected the C-strain mtCOI haplotype in Australia at very low frequencies. Proportions of C-strain to R-strain also varied significantly across the different SEA populations (Table S1) in contrast to the patterns observed in China, India, and African nations (e.g., Otim et al. 2018b, Nagoshi et al. 2020, Zhang et al. 2020, Jiang et al. 2022, Tay et al. 2022d). All Australian populations analysed for their corn or rice mitochondrial haplotypes via mitogenome assemblies of whole genome sequencing data therefore contrasted with the invasive populations from SEA where in some countries (e.g., Myanmar, Vietnam) FAW with the C-strain mtCOI haplotype made up approximately 50% of the populations examined (Table S1, Figs. 2a, 2b).

Mutations in the Triosephosphate isomerase (*Tpi*) gene used to differentiate the C- and R-strains suggested that virtually all individuals from the SEA, PA, and South Korean (i.e., EA) carried the C-strain genotype at the *Tpi* gene. Of all SEA and PA individuals assessed, none was homozygous for the rice allele at this locus, while a number of samples (n = 10) were heterozygous C-/R-strains, primarily from the Philippines as well as one sample each from Malaysia and South Korea. Similar to the conclusions of Zhang et al. (2020), at least in the invasive range, our finding suggested that the *Tpi* is not a useful marker: not only are the majority of samples of ‘corn’ preference, with limited evidence of hybrids and therefore contrasting greatly with whole genome analysis, but both the *Tpi* and the mitochondrial markers frequently contradicted each other, and do not provide accurate genealogy in, e.g., hybrid females or their offspring, especially if the female also mated with a hybrid male.

### Mitochondrial Genome analysis

In the invasive populations, we identified for the C-strain at least 12 unique maternal lineages forming a total of five clades. For the R-strain, conservatively we identified 19 maternal lineages from nine clades (i.e., I - IX; Fig. 1 ‘C-strain cladogram’). Combining results from this study and the study of Tay et al. (2022d), we therefore identified at least 27 unique maternal lineages in the invasive FAW populations from across 14 countries from Africa, South Asia, East Asia, SEA/Pacific, and Oceania.

**Fig. 1:**
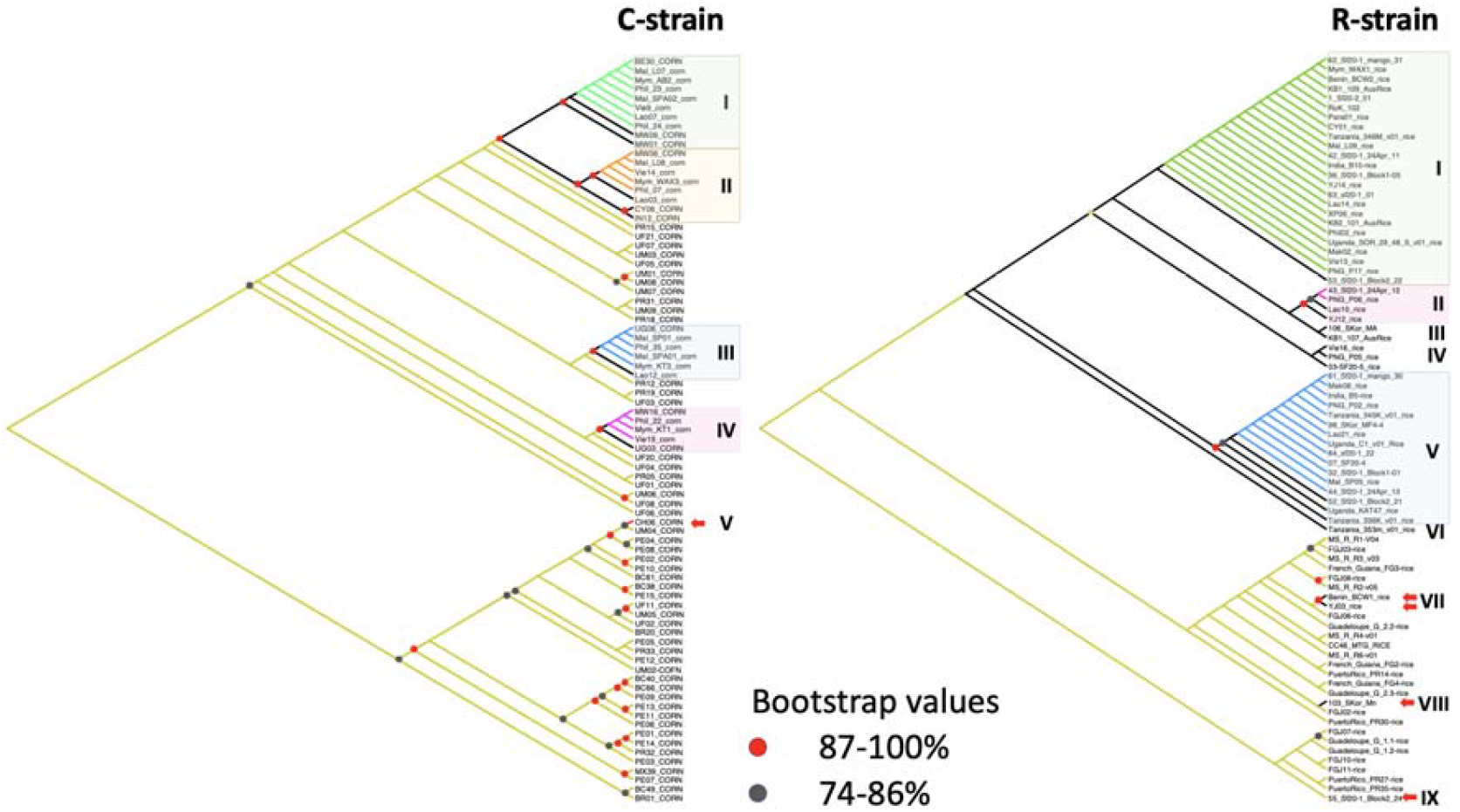
Maximum Likelihood cladograms of unique *Spodoptera frugiperda* C-strain and R-strain partial mitochondrial genomes based on concatenation of the 13 PCGs (11,393 bp) using IQ-Tree with 1,000 UFBoot replications. Individuals in clades I, II, III, and IV (C-strain) and in Clades I, II, V (R-strain) that are in the same colour scheme (i.e., green, orange, blue, or pinks) shared 100% nucleotide identity. Mitogenome haplotypes from native individuals for both C- and R-strains are in khaki green colour. Red and dark grey dots at branch nodes represent bootstrap values of 87-100% and 74-86%, respectively. Bootstrap values <74% are not shown. Red arrows indicate invasive individuals’ mitogenome haplotypes that are nested within native individuals. Country and sample codes are listed in Tables S1 and S2.

**Fig. 2:**
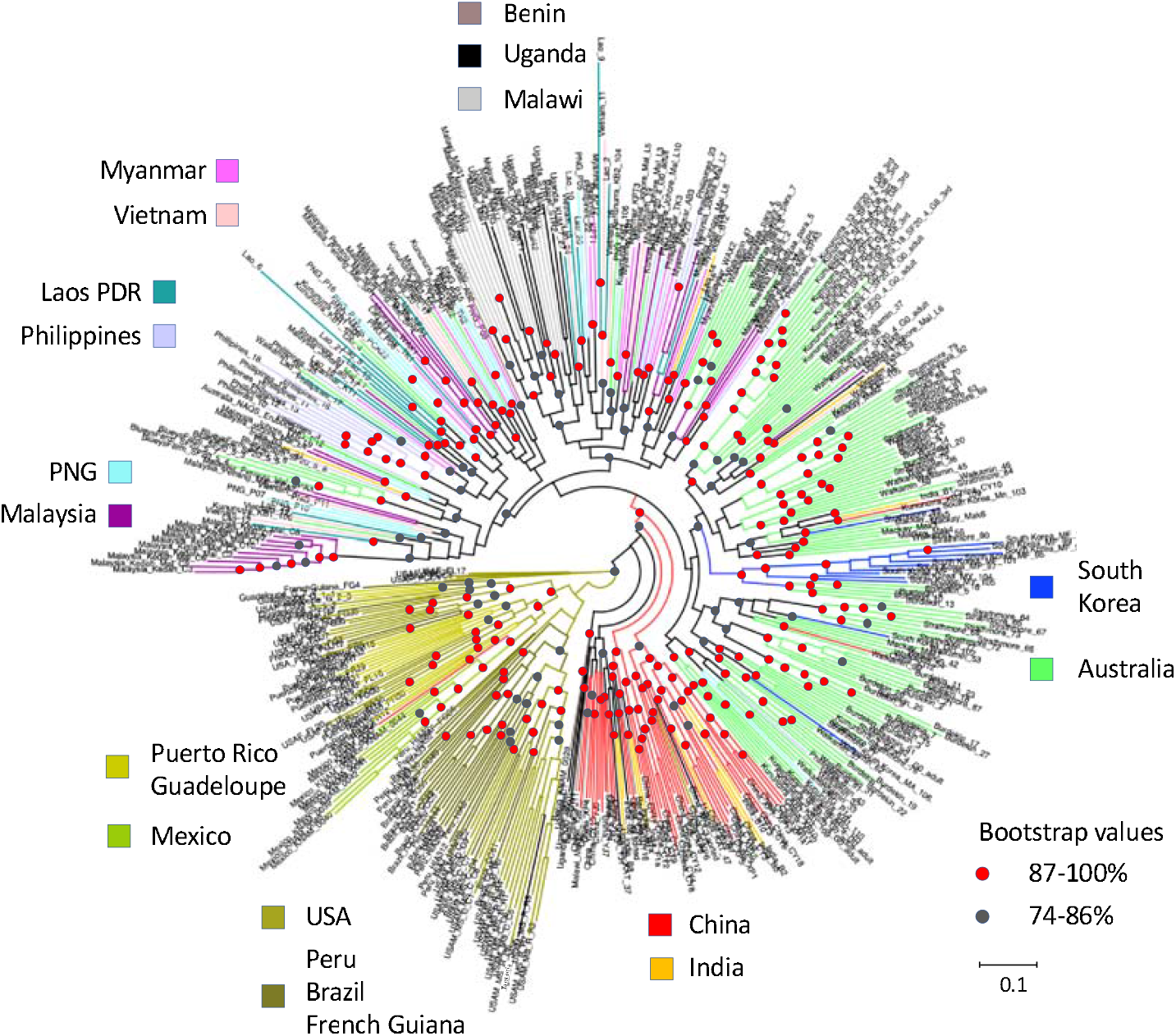
Maximum Likelihood (ML) phylogeny using 879 genome-wide non-coding SNPs with 1000 ultrafast (UF) bootstrap replications. Node support with <74% are not shown., 74-86% are represented by grey circles, 87-100% are represented by red circles. Country of origin is colour coded.

The C-strain mitochondrial genomes detected in Southeast Asia (Clade II) differed from the one present in the Yunnan CY and Indian populations although the four haplotypes clustered with high (87-100%) bootstrap support. One other C-strain mitogenome was detected in one individual from China (Tay et al. 2022d) and this remained a singleton haplotype amongst the other eight unique C-strain mitogenomes detected from African and SE Asian populations. For the R-strain mitogenomes (Fig. 1 ‘R-strain cladogram’), Clade I consisted of invasive individuals from all countries sampled to-date, although it did not contain individuals from all populations (e.g., no individuals from Australia Sf20-4, Sf20-5 or Mackay populations). R-strain Clade I also contained unique mitogenomes of individuals from the three China Yunnan populations, although other Chinese R-strain mitogenome haplotypes were also detected in clades II and VI. The second largest clade (Clade V) contained R-strain mitogenomes shared between individuals from six Australia populations, East Africa (Tanzania, Uganda), India, South Korea, Laos, PNG, and Malaysia (i.e., blue branches), although excluded other populations from SEA (i.e., Myanmar, Vietnam, Philippines), China, and Africa (i.e., Malawi, Benin). Clades II, III, IV, VIII, IX contained mitogenome haplotypes of individuals from either SEA and/or Asia (China, India) but not individuals from any of the African countries. Clades VII, VIII, and IX also have unique mitogenome haplotypes of invasive individuals that were more similar to French Guiana (FG code) or the Caribbean (i.e., Puerto Rico (PR), FG, or Guadeloupe) populations. The overall low bootstrap confidence at various nodes for both C-strain and R-strain phylogenies reflect the high number of maternal lineages with low mitogenome nucleotide diversity (i.e., mitogenome haplotypes differed by only very few nucleotide substitutions) across both invasive and native range populations. These limitations therefore lower their powers for use to confidently infer invasive population origins.

### Nuclear SNP Phylogenetic analysis

Maximum Likelihood (ML) phylogenetic analysis of the global *S. frugiperda* populations based on 870 neutral single nucleotide polymorphic (SNP) markers provided evidence of complex genetic relationship especially between individuals representing the invasive populations (Fig. 2). While the New World native populations (USA, Puerto Rico, Guadeloupe, Mexico, French Guiana, Peru, Brazil) clustered together and shared a basal (ancestral) lineage, the invasive populations did not identify Africa as the founding invasive population nor support the axiom of rapid west-to-east spread across the Old World. Selected individuals from Uganda and Benin appeared to cluster close to China/India populations, but there was no strong bootstrap support for a shared common ancestor (i.e., bootstrap value <74%). China populations (i.e., CY, YJ, XP from Yunnan; see Fig. 1B in Guan et al. 2020) formed two closely related sister clades with long branch lengths, suggesting these likely represented separate introduction events. The main Australian FAW population clade consisted of multiple sister clades and were clustered with varying degree of bootstrap support values. Some of these Australian populations appeared to shared close relationship with FE (i.e., South Korean) population (>87-100% support (e.g., selected Burdekin individuals)), while others had largely unknown invasive origin(s).

Southeast Asian countries (i.e., Malaysia, Philippines, Laos PDR, Vietnam, Myanmar) and Pacific/Australia (i.e., PNG) have populations that were mixed, and interestingly were ancestral to East African populations (i.e., Malawi, Uganda), while some Australian populations (i.e., NSW; NT) were shown to be closely related to these SE Asian populations with strong bootstrap support (e.g., >87-100% support for NT and NSW populations). Myanmar and China appeared to not share FAW populations with similar SNP profiles despite being neighbours. Multiple introductions of FAW with diverse genetic profiles and potentially similarly diverse origins were therefore likely the main reason for their wide spread detection across the Old World.

Clear evidence of multiple introductions can therefore be seen from the phylogenetic analysis, with unique introduction events likely underpinning the establishment of FAW populations in EA. (e.g., China) and SEA (e.g., Malaysia). In Australia, while the NT/NSW populations shared closer evolutionary relationships with SE Asian populations, the population origin for most of Qld and WA populations was unclear, and could be due to a sampling effect (e.g., this study could not source FAW populations from e.g., Indonesia, Cambodia, Thailand, Sri Lanka, Bangladesh, etc.). Some Burdekin and South Korean individuals appeared to share closer evolutionary lineage, although the factor leading to this detected pattern remained as yet unknown.

### Population Statistics and estimates of substructure

The basic population diversity statistics for each population are listed in Table 2. Nucleotide diversity (π) varied across a narrow range with the lowest (0.237) being from the Malaysian Kedah State population (i.e., reflected its lab-colony background), and the highest from the Malawian population (0.324), similar to that reported (Tay et al. 2022d; π = 0.279 – 0.329) in native and invasive populations based on the same set of 870 SNP loci. The high nucleotide diversity estimates from the SEA, Australia and South Korean populations likely reflected effects of limited (i.e., 870) highly polymorphic SNPs from non-coding genomic regions being used, and are comparable to the findings from Tay et al. (2022d) that used the same sets of SNP loci. We did not detect significant overall differences between the various invasive populations, and for the SE Asian and Australian populations the nucleotide diversity estimates were generally between 0.258-0.291. All invasive populations from Africa (Benin, Malawi, Uganda), SA/EA (India, China (CY, XP, XJ), South Korea), SEA (PNG, Malaysia (Johor, Kedah, Penang), Laos, Vietnam, Myanmar, Philippines), and Australia (WA (Kununurra), NT, QLD (Strathmore, Walkamin, Burdekin), NSW (Wee Waa)) showed higher average observed heterozygosity (Het_obs_) than the average expected heterozygosity (Het_exp_), with the highest Het_obs_ seen in the Malawian population as also reported previously (Tay et al. 2022d). Within the Australian populations, highest average observed heterozygosity was seen in the Strathmore population, while in populations in SEA (i.e., Myanmar, Philippines) and the EA (i.e., South Korea) all showed similar average Het_obs_. Negative average *F*_IS_ values for all populations were consistent with Het_obs_ being higher than Het_exp_, and suggested populations were out-breeding (i.e., avoidance of matings between related individuals; Wright 1965). Similar to the previous findings (Tay et al. 2022d), the lower Het_exp_ (i.e., Het_obs_ > Het_exp_; see Luikart and Cornuet 1998) could likewise indicate recent mixing of distinct populations from SEA that suggest multiple introductions (e.g., Jiang et al. 2022, Tay et al. 2022d *cf*. Cock et al. 2017, FAO 2018, Nagoshi et al. 2019b, Nagoshi et al. 2019a; i.e., due to a recent bottleneck from a recent western Africa founder event).

The observed heterozygosity excess detected in all invasive range populations could be further explained as due to population sub-structure and isolation breaking through periodic migration. Significant numbers of loci (*ca*. 30%) were also shown to not be in Hardy-Weinberg equilibrium (HWE) especially for the Malaysian (i.e., Kedah), but also Australian (i.e., Wee Waa, NT, Kununurra), Chinese (e.g., XP), South Korean, and Malawian populations. Taken as a whole, genetic diversity results from this study therefore suggested that the invasive Asian (i.e., SA, SEA, EA) FAW populations exhibited signatures of recent mixing of previously separated populations. Simulated patterns of moth migration of various invasive FAW populations such as between Myanmar and China (e.g., Sun et al. 2019, Wu Q et al. 2019, Li et al. 2020) and to Australia (Qi et al. 2021) are incompatible with the population genomic data which suggests these were likely discrete and non-panmictic FAW populations with the most probable explanation being due to multiple origins of founding populations.

### Genetic differentiation analysis

Estimates of pairwise genetic differentiation (*F*_ST_) between populations varied significantly (Table 2) and extended to between populations within a country (e.g., Mackay vs. rest of Australia; Kedah vs. rest of Malaysia). Of interest are the pairwise estimates between different Australian FAW populations from Kununurra (Western Australia), Northern Territory, Queensland (Strathmore, Walkamin, Burdekin, Mackay) and New South Wales (Wee Waa) that represented the most recently reported invasive populations in this study, and predominantly showed significant differentiation amongst themselves (with the exception of the two Queensland populations of Mackay and partially for Walkamin) and with other SEA/SA/EA countries. The majority of non-significant population genetic differentiation estimates were in SEA where the presence of FAW was reported earlier, i.e., since 2018 (e.g., FAO 2019, 2020) or as early as 2008 (Vu 2008, Nguyen and Vu 2009; see also Tay et al. 2022d), while across Asia (e.g., China) since 2016 but also potentially pre-2014 (Gilligan and Passoa 2014, Tay and Gordon 2019; see also Tay et al. 2022d).

Interestingly, significant genetic differentiation was observed between populations from Yunnan province in China and populations from Myanmar, Laos, and Vietnam. Penang and Johor (Malaysia) populations were not significantly differentiated from other SE Asian populations, nor with Ugandan and Malawian populations from east Africa. Individuals from Benin and Mackay (Queensland, Australia) showed non-significant genetic differentiation with all populations except with Kedah, and for Mackay also surprisingly with the Wee Waa population from New South Wales. The South Korean population exhibited significant genetic differentiation with SE Asian population except with Mackay, India and the Yuanjiang (YJ) population in Yunnan Province. Finally, the Kedah population, being one of the earliest collected samples from Malaysia and having been maintained as a laboratory population, showed strong differentiation with all populations (and lowest nucleotide diversity, π = 0.237; Table 1) further supporting unique, non-African, introduction events in SEA. Strong genetic differentiation suggested there was limited gene flow to breakdown sub-structure between populations, and the *F*_ST_ estimates from these invasive populations therefore failed to support a west-to-east spread pathway for the FAW. This observation instead suggested the widespread presence of genetically distinct FAW populations, likely due to independent introductions and therefore also highlighting likely biosecurity weaknesses especially in East Asia (e.g., China, South Korea) and SEA (e.g., Malaysia).

**Table 1:** Population statistics for *Spodoptera frugiperda* populations from Southeast Asia (i.e., Malaysia (MYS; Johor, Kedah, Penang), Laos, Vietnam, Myanmar), East Asia (i.e., South Korea), and Pacific/Australia (i.e., Papua New Guinea (PNG), Australia). Populations from Australia (AUS) are Kununurra from Western Australia (WA), Northern Territory (NT), from Queensland (Qld; Strathmore, Walkamin, Burdekin, Mackay), and New South Wales (NSW; Wee Waa). Invasive populations from Benin (BE), Uganda (UG), Tanzania (TZ), Malawi (MW), India (IN), and China (CH, CC, CY, CX) were from (Tay et al. 2022d). Nucleotide diversity: (Nt Div, π); Avg. Het_exp_: average expected heterozygosity; Avg. Het_obs_: average observed heterozygosity; Avg. Hom_exp_: average expected homozygosity; Avg. Homt_obs_: average observed homozygosity; % HWE: per centage loci in Hardy-Weinberg Equilibrium; F_IS_: inbreeding coefficient. Refer to Table S1 and Methods sections for population details and how the statistics were calculated.

| PopID | % HWE | Av. Het (Obs) | s.e. | Av. Hom(Obs) | s.e. | Av. Het(Exp) | s.e. | Av. Hom(Exp) | s.e. | Nt. Div ($\pi$) | s.e. | Av. Fis | s.e. |
| --- | --- | --- | --- | --- | --- | --- | --- | --- | --- | --- | --- | --- | --- |
| AUS_Kununurra | 0.69 | 0.386 | 0.011 | 0.614 | 0.011 | 0.272 | 0.006 | 0.728 | 0.006 | 0.277 | 0.006 | -0.265 | 0.075 |
| AUS_NT | 0.62 | 0.381 | 0.015 | 0.619 | 0.015 | 0.236 | 0.008 | 0.764 | 0.008 | 0.258 | 0.009 | -0.239 | 0.012 |
| AUS_Strathmore | 0.70 | 0.407 | 0.011 | 0.594 | 0.011 | 0.286 | 0.006 | 0.714 | 0.006 | 0.291 | 0.006 | -0.277 | 0.040 |
| AUS_Walkamin | 0.74 | 0.394 | 0.010 | 0.606 | 0.010 | 0.281 | 0.006 | 0.719 | 0.006 | 0.286 | 0.006 | -0.264 | 0.038 |
| AUS_Burdekin | 0.70 | 0.395 | 0.011 | 0.605 | 0.011 | 0.281 | 0.006 | 0.719 | 0.006 | 0.286 | 0.006 | -0.266 | 0.037 |
| AUS_Mackay | 0.72 | 0.391 | 0.012 | 0.609 | 0.012 | 0.269 | 0.007 | 0.731 | 0.007 | 0.290 | 0.007 | -0.212 | 0.013 |
| AUS_Wee Waa | 0.64 | 0.393 | 0.013 | 0.607 | 0.013 | 0.257 | 0.007 | 0.743 | 0.007 | 0.275 | 0.008 | -0.245 | 0.017 |
| PNG | 0.72 | 0.403 | 0.011 | 0.597 | 0.011 | 0.284 | 0.006 | 0.716 | 0.006 | 0.292 | 0.006 | -0.261 | 0.022 |
| Johore_MYS | 0.80 | 0.400 | 0.011 | 0.600 | 0.011 | 0.279 | 0.006 | 0.721 | 0.006 | 0.294 | 0.006 | -0.238 | 0.017 |
| Kedah_MYS | 0.35 | 0.377 | 0.016 | 0.623 | 0.016 | 0.224 | 0.009 | 0.776 | 0.009 | 0.237 | 0.009 | -0.281 | 0.030 |
| Penang_MYS | 0.75 | 0.396 | 0.012 | 0.604 | 0.012 | 0.271 | 0.007 | 0.729 | 0.007 | 0.287 | 0.007 | -0.236 | 0.015 |
| Laos | 0.79 | 0.390 | 0.010 | 0.610 | 0.010 | 0.282 | 0.005 | 0.718 | 0.005 | 0.289 | 0.006 | -0.246 | 0.034 |
| Vietnam | 0.87 | 0.404 | 0.011 | 0.596 | 0.011 | 0.286 | 0.006 | 0.714 | 0.006 | 0.300 | 0.006 | -0.238 | 0.020 |
| Myanmar | 0.73 | 0.410 | 0.011 | 0.590 | 0.011 | 0.289 | 0.006 | 0.711 | 0.006 | 0.298 | 0.006 | -0.264 | 0.017 |
| India | 0.77 | 0.406 | 0.010 | 0.594 | 0.010 | 0.289 | 0.006 | 0.711 | 0.006 | 0.301 | 0.006 | -0.243 | 0.000 |
| ChinaCY | 0.73 | 0.399 | 0.011 | 0.601 | 0.011 | 0.279 | 0.006 | 0.721 | 0.006 | 0.287 | 0.006 | -0.263 | 0.000 |
| ChinaXP | 0.68 | 0.404 | 0.011 | 0.596 | 0.011 | 0.280 | 0.006 | 0.720 | 0.006 | 0.290 | 0.006 | -0.262 | 0.000 |
| ChinaYJ | 0.71 | 0.403 | 0.011 | 0.597 | 0.011 | 0.280 | 0.006 | 0.720 | 0.006 | 0.292 | 0.006 | -0.251 | 0.000 |
| Philippines | 0.74 | 0.412 | 0.011 | 0.588 | 0.011 | 0.289 | 0.006 | 0.711 | 0.006 | 0.297 | 0.006 | -0.271 | 0.026 |
| Skorea | 0.68 | 0.413 | 0.012 | 0.587 | 0.012 | 0.278 | 0.007 | 0.722 | 0.007 | 0.291 | 0.007 | -0.267 | 0.023 |
| Uganda | 0.75 | 0.434 | 0.010 | 0.566 | 0.010 | 0.306 | 0.005 | 0.694 | 0.005 | 0.316 | 0.005 | -0.275 | 0.000 |
| Malawi | 0.69 | 0.449 | 0.010 | 0.551 | 0.010 | 0.314 | 0.005 | 0.686 | 0.005 | 0.324 | 0.005 | -0.291 | 0.000 |
| Benin | 0.79 | 0.415 | 0.013 | 0.586 | 0.013 | 0.277 | 0.007 | 0.723 | 0.007 | 0.317 | 0.008 | -0.184 | 0.000 |

**Table 2:**
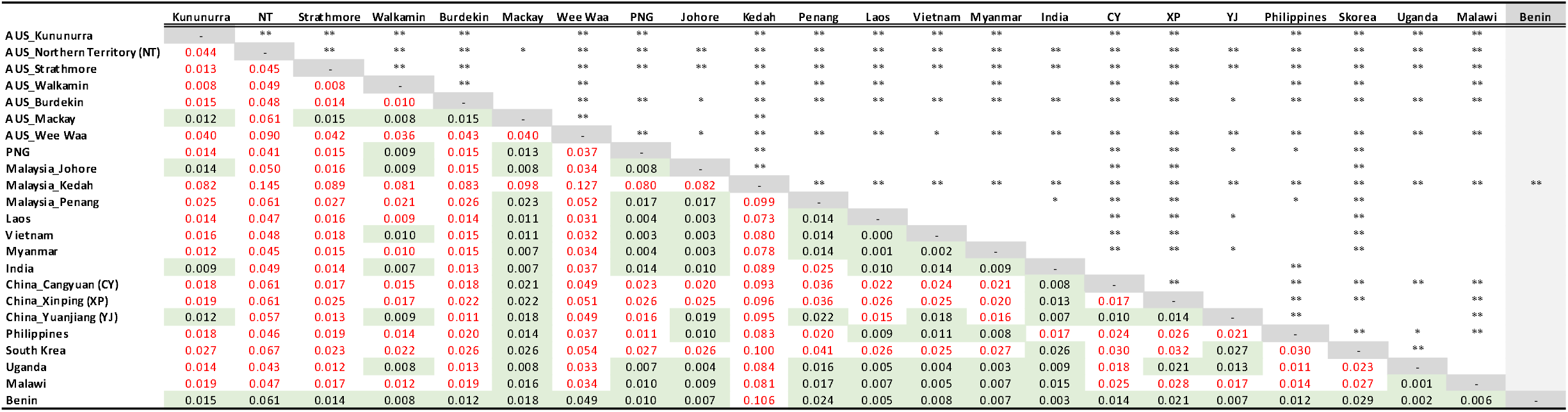
Population genetic differentiation via pairwise *F*_ST_ estimates between *Spodoptera frugiperda* populations from the invasive ranges of Africa (Uganda, Malawi, Benin), South Asia (India), East Asia (China, South Korea), Southeast Asia (Malaysia (Johor, Kedah, Penang States), Laos, Vietnam, Myanmar), and Pacific/Australia (Papua New Guinea (PNG), AUS; Kununurra (Western Australia, WA), Northern Territory (NT), Strathmore, Walkamin, Burdekin, Mackay (Queensland, Qld), Wee Waa (New South Wales, NSW). Estimates of significance are: (*) P ≤ 0.05; P ≤ 0.01 (**). Values in red are significant differentiation between pairwise comparisons, values in green cells are not significant. Western African (Benin) population (light grey column) showed non-significant pairwise F_ST_ values with all locations except with the Malaysian Kedah State population. Refer to Table S1 and Methods section for population details and statistical calculation approach. Refer to (Tay et al. 2022d) for details of African and Asian FAW populations.

**Table 3:**
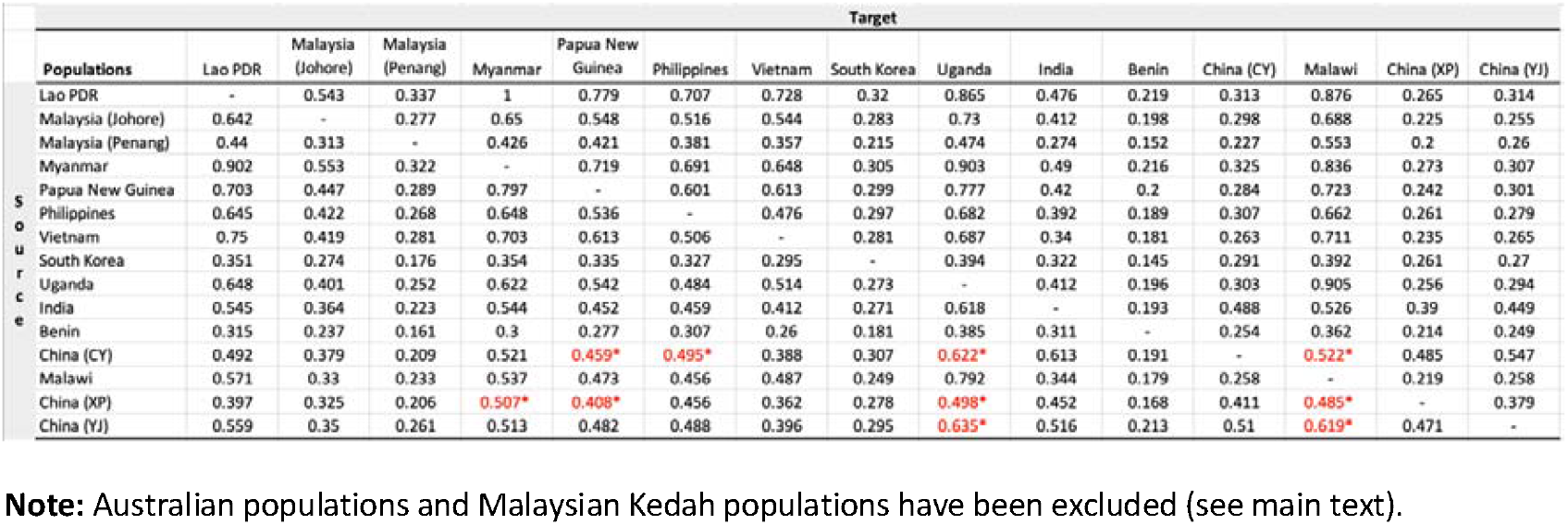
DivMigrate matrix showing effective migration rates calculated using GST from source to target invasive populations. Significant unidirectional migration detected from 100 bootstrap replications (at alpha = 0.5) are in red and indicated with *. All three Chinese populations were from the Yunnan province (CY: Cangyuan, XP: Xinping, YJ: Yuanjiang). Malaysian populations are from the states of Johor and Penang as indicated in parentheses.

The genetic diversity of Australian populations identified surprisingly complex sub-structure patterns given the short time frame of population detections across different northern Australian regions. Significant genetic differentiation between, e.g., Kununurra (WA), Northern Territory (NT), Queensland (e.g., Strathmore, Burdekin), and Wee Waa (NSW) populations suggests these populations likely derived from separate establishment events. The WA Kununurra population was not significantly differentiated from the Johor State (Malaysia), India and the Cangyuan (CY) China populations, suggesting a potential south-eastern route from SA/SEA into north-western Australia. Contrasting this, Walkamin and Mackay populations showed non-significant genetic differentiation with the Madang (PNG) population, suggesting a potential second pathway for SEA individuals to arrive into the north-eastern region of Australia. Significant genetic differentiation between WA, NT, and Qld populations suggested that at least during the early stage of pest establishment in northern Australia, there was limited gene flow to harmonise the unique genetic background carried by these distinct individuals, some of which exhibited also distinct insecticide resistance profiles (Tay et al. 2022b).

### PCA

We selected specific populations to compare using Principal Component Analysis (PCA) as examples to support evidence of independent introductions, as seen from Fig. 3a between China (CY, YJ, XP) populations vs. Myanmar, in Fig. 3b (within Malaysian populations between those collected from Penang and Johor States vs. Kedah State), in Fig. 3c for between China and East Africa (e.g., Uganda, Malawi), and where Benin and India individuals that grouped with either China or east Africa; and in Fig. 3d between China, Malaysia (Kedah State), and Australia (NT, NSW)). Genetic variability between Australian populations (e.g., Strathmore (QLD) vs. NT and NSW) was also evident (Fig. 3d).

**Fig. 3:**
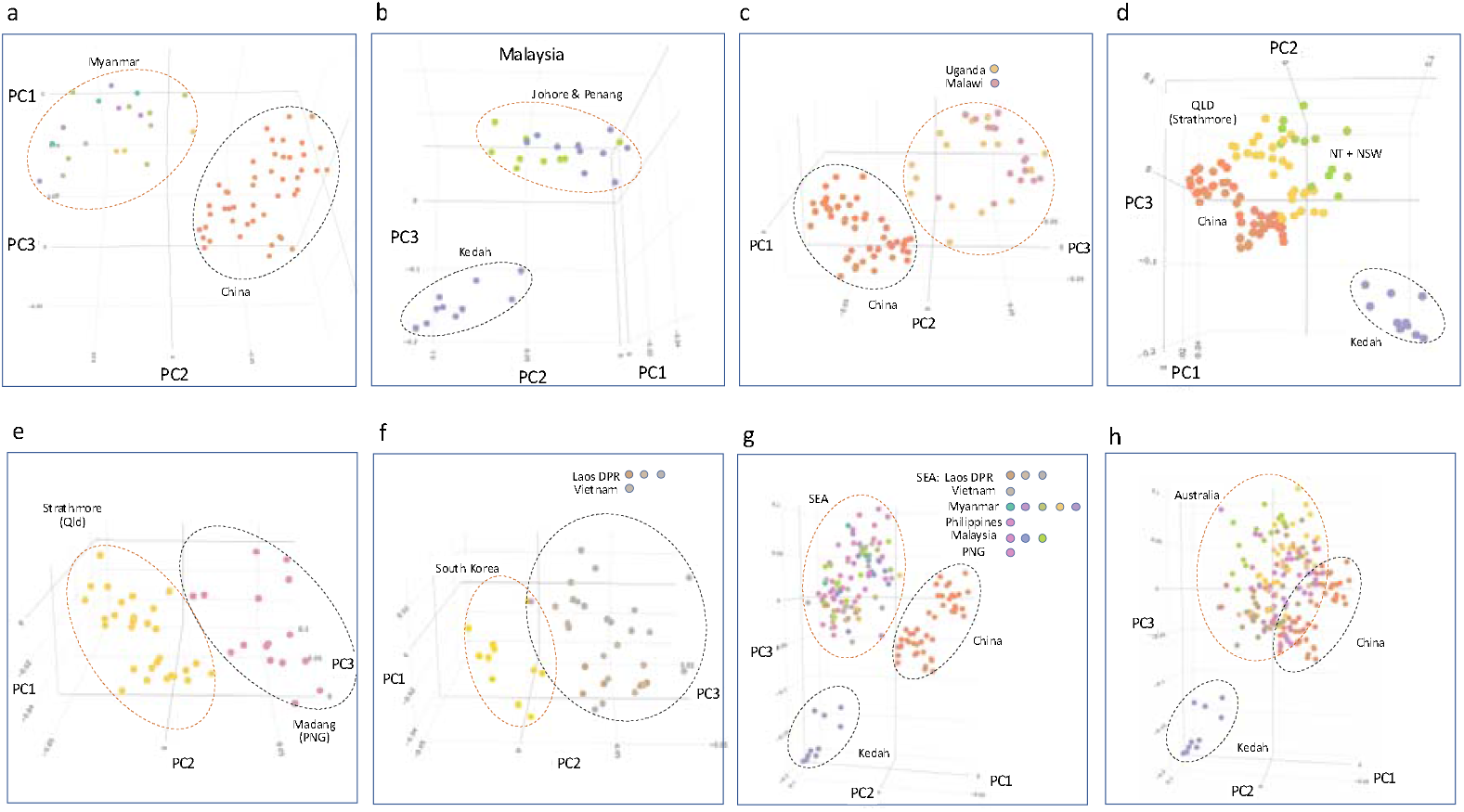
Principal component analysis (PCA) showing variability between selected FAW populations from their invasive ranges. **(a)** China and Myanmar; **(b)** Kedah and Johor/Penang populations from Malaysia, **(c)** China and east African (Uganda/Malawi) populations, **(d)** Australia (Strathmore, Qld/Northern Territory + New South Wales), China, and Malaysia (Kedah) populations, **(e)** Australia (Strathmore, Qld) and PNG (Madang Province) populations, **(f)** Lao PDR/Vietnam and South Korea populations, **(g)** China and SE Asian (Lao PDR/Vietnam/Myanmar/Philippines/Malaysia) and Pacific/Australia (PNG) populations, and **(h)** Australia, China and Malaysia (Kedah) populations. Note the overall population genomic variability between countries (e.g., a, **c-g**) and within countries (e.g., Malaysia **(b)**, Australia **(d)**). Populations with similar genomic variability are also evident, e.g., for Strathmore **(e)** and South Korea **(f)**; and for Madang **(e)** and Lao PDR/Vietnam **(f)**, further supporting potential different population origins of various FAW populations across the current invasive regions. The Southeast Asian and Chinese populations are overall different **(g)**, Australia’s FAW populations showed similarity with both Southeast Asia and China **(g, h)**.

PCA also showed that differences existed between FAW populations from the Madang Province in PNG and with the Strathmore population from Qld (Fig. 3e). The SEA FAW populations from Lao PDR/Vietnam also exhibited diversity from the South Korean population (Fig. 3f), with the South Korean and Strathmore populations largely exhibiting similar diversity pattern, while the Madang population shared similarity with Laos and Vietnam populations. Plotting all SEA populations against China clearly showed that populations from SEA were distinct from the Chinese FAW populations (Fig. 3g), while in Australia individuals from various populations shared similarity with both Chinese and SEA FAW. Despite the connectedness of the landscape between SEA and China, SEA largely appeared to have their own FAW populations, with FAW in SEA and in China differing in their genome composition overall as shown via PCA.

PCA further enabled visualisation of genetic diversity amongst Australia FAW populations, suggesting that arrival and establishment of FAW likely involved separate introduction events that followed closely after each other and over a short timeframe. While it had been anticipated that the southward spread of FAW from SEA would necessarily lead to Australian FAW and PNG FAW to share similar genetic backgrounds, the Madang Province FAW population appeared to be different from the Strathmore (Qld) population, with the Madang population being more similar to Lao PDR/Vietnam populations, and the Strathmore population more similar to FAW from South Korea.

### DivMigrate analysis

Directionality of gene flow between African, South Asia (Indian), East Asia (China) and SE Asian populations were predominantly from China to east African and Southeast Asian populations (e.g., Figs. 4a, 4b, Fig. S-1), while movements of FAW in Laos and Viet Nam (i.e., the Indochina region) were predominantly with other SEA countries (e.g., with Myanmar and East Africa; Figs. 4c, 4d, Fig. S-2) but with no directional movements to the three Yunnan populations (CY, XP, YJ). Migration directionality with other SE Asian populations (e.g., Johor (JB; Fig. S-3) and Penang (PN, Fig. S-4)) showed that these two populations (but especially the Johor population) were predominantly source populations for Uganda, Malawi, Philippines, Vietnam and PNG (Fig. S-3). Bidirectional migration between Myanmar and Laos PDR populations were also detected with the Johor population from Malaysia (Fig. S-3). When India was selected as the source population, bidirectional migration events were detected with Myanmar and with the Cangyuan (CY) populations (Fig. S-5) while unidirectional migration events from India to Uganda and Malawi and to Laos were detected, and the China Yuanjian (YJ) population showed unidirectional migration to India. Unidirectional migration events from CY and YJ populations to the PNG Madang population were detected, while bidirectional migration events between PNG and Myanmar, Laos PDR, Philippines, Vietnam, and with Uganda and Malawi were also detected (Fig. S-6). No migration events were detected between the West African Benin population and with the South Korean population.

**Figs. a, 4b:**
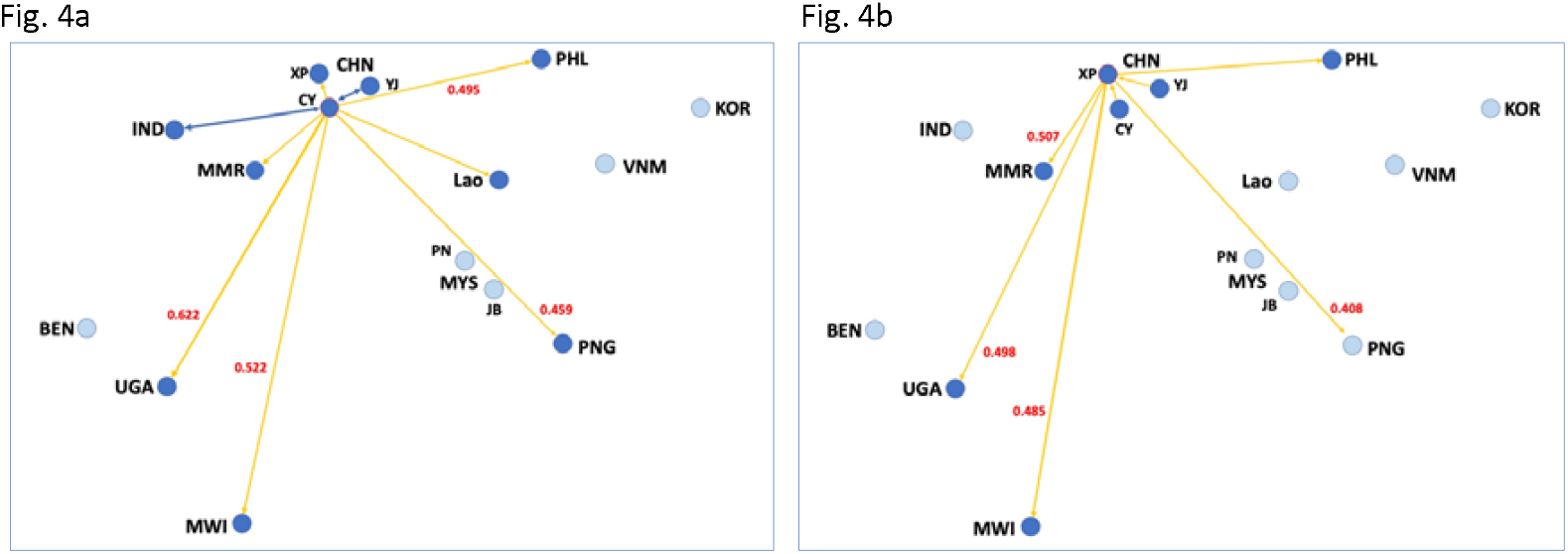
DivMigrate analyses with edge weight setting at 0.453 showing unidirectional (yellow arrow lines) and bidirectional (blue arrow lines) gene flow between countries in Africa and South Asia/East Asia/SE Asia. Significant migration rates (at alpha = 0.5) are in red and as provided in Table 3. Incidences of unidirectional migration were predominantly detected from China (CHN) Yunnan populations (CY, XP) to SE Asian populations (e.g., Myanmar (MMR), Laos PDR (Lao), Philippines (PHL)) and to east African populations (e.g., Uganda (UGA), Malawi (MWI)) (Fig. 4a, 4b). Source populations are CY (Fig. 4a) and XP (Fig. 4b).

**Figs. 4c, 4d:**
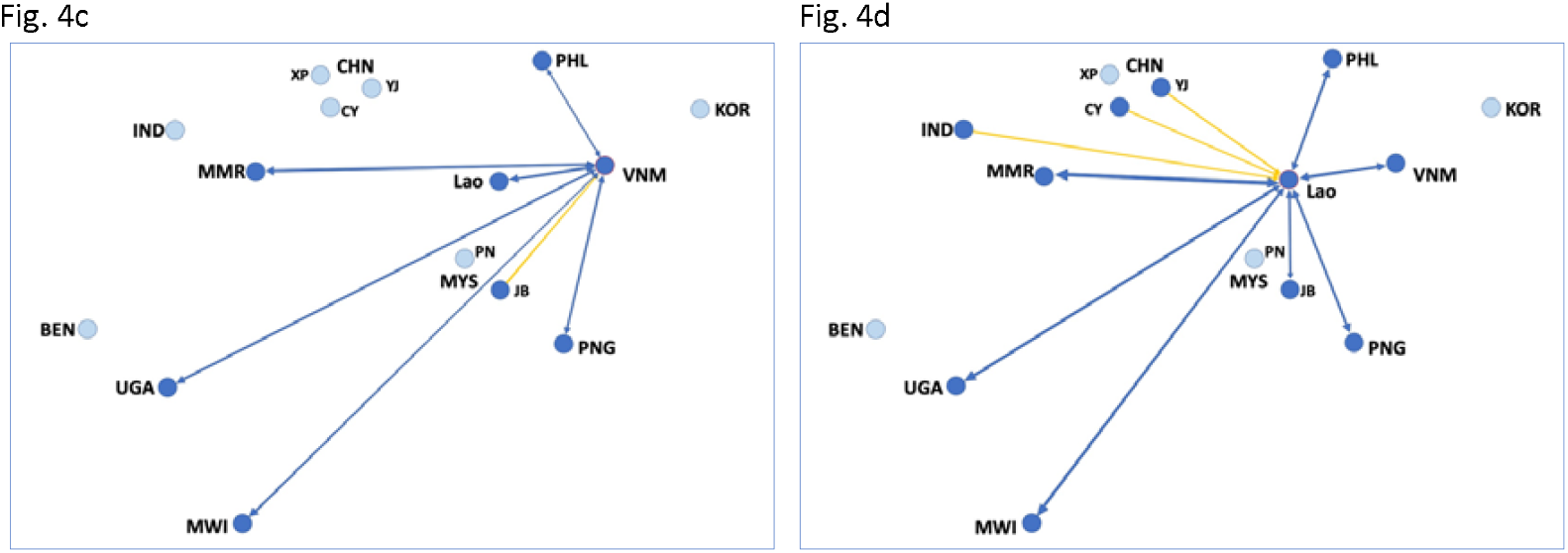
DivMigrate analyses with edge weight setting at 0.453 showing unidirectional (yellow arrow lines) and bidirectional (blue arrow lines) migration between countries in Africa and South Asia/East Asia/SE Asia. Migration rates between populations are as provided in Table 3. **Fig. 4c:** Vietnam (VNM) as the source population identified an incidence of unidirectional migration from Malaysia (MYS) Johor state (JB) to Vietnam, while bidirectional migration events were detected from Vietnam to other SE Asian (e.g., Philippines (PHL), Lao PDR (Lao), Myanmar (MMR)), to Pacific/Australia (i.e., Papua New Guinea (PNG)), as well as to east Africa (Uganda (UGA), Malawi (MWI)). **Fig. 4d:** Lao PDR (LAO) as source population identified bidirectional migration events between various SEA populations and east African populations, while unidirectional migration events were identified from India (IND) and China (CHN) Yunnan populations (CY, YJ) to Laos PDR. No migration events were evident from SE Asian populations to China.

### Admixture analysis

Admixture analyses involving all Australian, Southeast Asian and South Korean populations from this study; and native populations from the Americas and Caribbean Islands, and invasive populations from Africa (Benin, Uganda, Malawi), India, and China (Tay et al. 2022d), provided an overall complex picture of population structure that reflected the species’ likely introduction histories across its invasive ranges.

Admixture analysis that excluded New World, African and Indian populations identified four genetic clusters (i.e., K = 4) to best describe these invasive populations from SEA, and EA (i.e., China, South Korea), and Pacific/Australia (Fig. 5a). At K = 4, Australian populations from NT and NSW, YJ population from China, South Korean, and Malaysia’s Kedah population, each showed unique admixture patterns (i.e., some individuals from NT and NSW populations lacked cluster 3; most of YJ (but also some CY and XP) individuals lacked clusters 1 and 2; South Korean (e.g., MF individuals) lacked cluster 2; Malaysia’s Kedah population lacked evidence of admixture (i.e., reflecting its laboratory culture history) and was made up predominantly by individuals that belonged to cluster 4. Populations from China also differed from most populations from SEA due to the overall absence of genetic cluster 4. Taken as a whole, establishment of the FAW populations in China, Malaysia, vs. other SE Asian populations, and between Australian populations (e.g., NT/NSW *cf*. WA/Qld), likely involved individuals from diverse genetic background (i.e., multiple introductions). At K = 4, the majority of Australian populations appeared to contain genetic clusters similar to China (i.e., cluster 3) and to SEA (i.e., cluster 2).

**Fig. 5:**
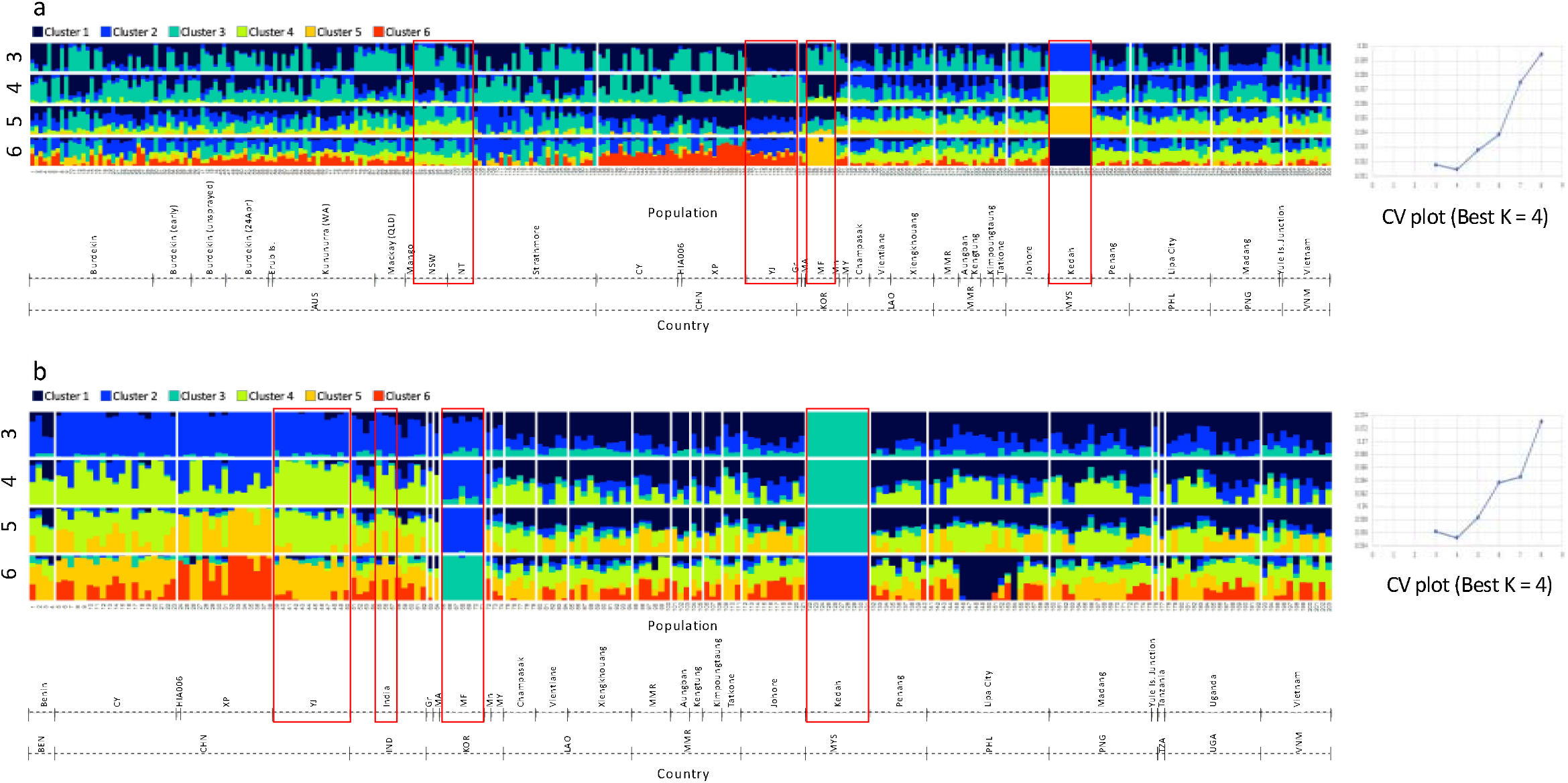
Admixture and corresponding CV plots for FAW populations from: **(a)** Australia, China, South Korea, Lao PDR, Myanmar, Malaysia, Philippines, PNG, and Vietnam, and **(b)** Benin, China, India, South Korea, Lao PDR, Myanmar, Malaysia, Philippines, PNG, Tanzania, and Vietnam. Optimal ancestral genetic clusters are K=4 for both admixture plots. Boxed individuals have unique admixture patterns at K=4 when compared with other populations. China FAW lacked Cluster 2 (navy blue colour; present in almost all SEA and Australian FAW), while in NSW and NT some individuals lacked cluster 3. South Korea ‘MF’ population generally lacked cluster 2, while Kedah (Malaysia) showed distinct (cluster 4) pattern for all individuals. The overall same observations are evident in the admixture plot in (b), with African FAW generally exhibiting admixture patterns similar to SEA populations than to Chinese FAW. With the exception of Kedah (Malaysia) and some Chinese FAW individuals, all FAW in the invasive range showed evidence of genomic admixture (i.e., hybrid signature).

Overall admixture patterns at best K = 4 in China and SEA remained unchanged when analysed together with African and Indian individuals (Fig. 5b; excluded Australia). Benin individuals were either similar to China or to SEA, while eastern African populations (e.g., Uganda, Malawi) were similar to Southeast Asian populations from e.g., Vietnam, Laos, and is in agreement with the phylogenetic inference (Fig, 3) that identified these African individuals as having loci that were derived from Southeast Asian populations.

Genome-wide SNP loci demonstrated that invasive FAW populations from SEA and Australia exhibited admixed genomic signatures similar to that observed in other invasive populations (Zhang et al. 2020, Tay et al. 2022d). While the current invasive populations in Africa and Asia likely arrived already as hybrids as suggested by Yainna et al. (2020), the Malaysia Kedah State population was potentially established by offspring of a non-admixed female. Distinct admixture patterns in Malaysian FAW populations between Kedah and Johor/Penang states therefore suggested that establishment of these populations was likely as separate introduction events. As reported also in Tay et al. (2022d), the Chinese YJ population appeared to have admixed signature that differed from the XP and CY populations, and suggested that the YJ population could have a different introduction history than the XP and CY populations. Similar multiple genetic signatures based on lesser nuclear markers by Jiang et al. (2022) also supported likely multiple introductions of China Yunnan populations.

## Discussion

In this study, we demonstrated through genomic analysis of genome-wide SNP markers signatures of multiple independent introductions of *S. frugiperda* in Asia (i.e., SEA. EA), the detection of distinct population sub-structures between what had been widely assumed (e.g., Sun et al. 2019, Wu Q et al. 2019, Li et al. 2020) as genetically linked populations between Myanmar and China, of multiple introduction pathways into north western and north eastern Australia, and the identification of potential SEA origins of eastern African (i.e., Uganda, Malawian) FAW populations. The findings highlight the need to infer population movements through genome-wide analyses; while critical assessments are needed of conclusions based on movement studies such as reverse trajectory analyses (Wu Q et al. 2019, Qi et al. 2021) and inference of population genetics relying solely on detection reports and limited (e.g., partial mitochondrial genes; partial TPI gene) DNA markers. Importantly, the study demonstrated the unexpected complexity of FAW populations across its current sub-Saharan, South Asia, Southeast Asia, East Asia, and Pacific/Australia invasive ranges since the species’ initial 2016 confirmation in western Africa, and were likely underpinned by human (anthropogenic)-assisted activities, as also cautioned by Early et al. (2018) based on the prevailing opposite wind direction (i.e., east-to-west) in the African continent.

The rapid spread rate of this pest since 2016 to over 70 countries (Czepak et al. 2019) has generally been regarded as a result of its long distance migration ability (e.g., Rose et al. 1975, Jones et al. 2019, Gui et al. 2020, Nagoshi et al. 2020) following a single introduction into west Africa (Goergen et al. 2016, Cock et al. 2017, Nagoshi et al. 2018, Nagoshi et al. 2019b, Nagoshi et al. 2019a) and subsequent wind-assisted dispersal (Day et al. 2017). In contrast, genomic (e.g., Schlum et al. 2021, Tay et al. 2022d) and genetic (Jiang et al. 2022, Tay et al. 2022c) evidence has begun to emerge that the current widespread establishments of FAW populations in Asia were the result of multiple introductions, and likely associated with international agricultural trade. While pre-border interception data from SEA are generally lacking, regular interceptions of FAW in agricultural commodities by Europhyte (e.g., <u>2018</u>) provided evidence to demonstrate the regular interceptions of FAW from its native countries (and increasingly, also from established invasive populations in Africa) were associated with international trade activities. These ‘bridgehead invasive populations’ (Guillemaud et al. 2011) established across Asia and Africa enabled subsequent establishments of localised secondary invasive populations via multiple factors including agricultural trade activities and natural migration (e.g., Early et al. 2018, Huesing et al. 2018, Jones et al. 2019).

From the analyses presented here, we suggest that the number of independent introductions of FAW is likely underestimated. For example, while our genomic analyses have identified populations in Malaysia, South Korea, and China (Tay and Gordon 2019, Tay et al. 2022d) with distinct signatures, Jiang et al. (2022) identified six separate genetic signatures in populations from southern and northern China, and Schlum et al. (2021) and Tay et al. (2022d) identified a total of least four separate introductions in African populations (e.g., Kenya, Benin, Uganda, Malawi). Population genetic analyses based on limited partial gene markers have also identified low frequencies of unusual polymorphism patterns in the *Tpi* and mtDNA gene markers in some African and an Indian FAW (e.g., Nagoshi et al. 2019b, Nagoshi et al. 2019a, Jing et al. 2021, Overton et al. 2021). Similarly, Gui et al. (2020) also detected signatures suggestive of independent introductions based on genome wide SNP analyses but have nevertheless maintained the African origin theory. While the conclusions by these authors have reinforced the hypothesis of the west-east movement of FAW, increasing whole genome evidence makes it more reasonable to conclude that these earlier single gene findings were also evidence of multiple unrecognised independent introductions.

Our investigation based on previously described genome-wide neutral SNP markers (Tay et al. 2021b) enabled integration of the published dataset (i.e., of native New World FAW populations and invasive African, Indian, and Chinese FAW populations) with the current SE Asia, East Asia, and Pacific/Australia FAW populations to reveal a complex picture of the pest’s invasive history. While multiple introductions especially in the Asian continent (e.g., China, Vietnam, Malaysia; Vu 2008, Nguyen and Vu 2009, Tay and Gordon 2019, Gui et al. 2020, Tay et al. 2021b, Jiang et al. 2022, Tay et al. 2022d; this study) have played an important role leading to the perception of rapid spread, introductions from elsewhere have also contributed to the African establishment and expansion. Further to genomics evidence, resistance allele characterisation and bioassay experiments have also identified invasive populations with non-overlapping unique insecticide resistance profiles in China (e.g., Lv et al. 2021), Indonesia (Boaventura et al. 2020), Australia (Tay et al. 2021a, Tay et al. 2022c), India (Deshmukh et al. 2020), and Africa (Eriksson 2019, Worku and Ebabuye 2019).

Multiple independent introductions of invasive agricultural pests are well-documented such as for the cryptic whitefly *Bemisia tabaci* MED and MEAM1 species (e.g., De Barro et al. 2011, de Moraes et al. 2018, Elfekih et al. 2018), and for the related noctuid pest *Helicoverpa armigera* (e.g., Tay et al. 2017, Arnemann et al. 2019, see also Jones et al. 2019), and underpinned the perceived rapid spread (e.g., Leite et al. 2014) of these pests in their introduced habitats. Early confirmation of invasive pests in novel habitats is also often challenging especially if initial population sizes were small. For example, while the initial detections of *H. armigera* in Brazil was during the cropping seasons of 2012/2013 (Czepak et al. 2013, Tay et al. 2013), its presence in Brazil was only subsequently confirmed to be at least as early as in 2008 (Sosa-Gomez et al. 2016) and potentially earlier elsewhere (e.g., in Peru) in the South American continent (Collins et al. 2007, Tay et al. 2017). Similarly for the FAW, its presence in Vietnam was reported since 2008 (Vu 2008, Nguyen and Vu 2009, Nguyen et al. 2012), and pre-border interceptions of FAW from non-native ranges (e.g., China, Indonesia, Israel, Micronesia, Netherlands, Thailand, and Turkey) back to its native range by the USDA Identification Technology Program (ITP) were pre-2014 (Gilligan and Passoa 2014, see also Tay et al. 2022d), and pre-dated the 2016 first reported cases of FAW in east Africa (Goergen et al. 2016). In Uganda, farmer surveys (Kalyebi 2020, Kalyebi et al. 2022) showed that FAW damage symptoms on maize crops were first identified in the Namutumba (since 2013) and the Kamuli (since 2014) districts, while (Otim et al. 2018a) reported farmers observed FAW damage symptoms in eastern and northern Uganda since 2014, although farmers at the time were unaware of this highly invasive exotic armyworm species. Findings of field surveys by Otim et al. (2018a), Kalyebi (2020), and Kalyebi et al. (2022) were in agreement with anecdotal observation and with single-gene response on Bt maize in South and East Africa, that the FAW could have been present in Africa several years prior to its confirmation in west Africa (Huesing et al. 2018).

While the current FAW populations in Vietnam were shown to have a gene flow directionality from Malaysia, whether this could be linked to the 2008 report (Vu 2008, Nguyen and Vu 2009) of outbreaks around Hanoi remained to be answered. Malaysian populations from Penang and Johor States were also shown to be linked to, and potentially source populations for, Ugandan and Malawian FAW populations (Fig. 2; Fig. S3; Fig. S4), in addition to the earlier results that also showed Chinese FAW populations to be potential source populations (Tay et al. 2022d; also Fig. 4a, Fig. 4b). Our study and that of Jiang et al. (2022) therefore identified two potential biosecurity weakness hotspots in Asia that have contributed to the spread of FAW in recent times. The study of Jiang et al. (2022) as well as the studies of Tay et al. (2022d) and Gui et al. (2020) therefore identified at least eight unique genetic/genomic signatures of introduction events across different regions (e.g., southern and northern regions) in China. While reverse trajectory studies by, e.g., Wu Q et al. (2019) suggested Yunnan province FAW to have originated from Myanmar, results from the present study showed that there was very little genetic connectivity between them (e.g., Fig. 3a), while DivMigrate analysis showed that gene flow directionality was more likely to be from Cangyuan (CY) to Myanmar (Fig. 4a), thereby highlighting the importance of incorporating genomic analyses to aid interpretations of pest movements.

Southeast Asian and Pacific/Australia countries such as Laos, Myanmar, Vietnam, Philippines, Malaysia, and PNG, appeared to be the geographic ‘genetic melting pot’ for the invasive FAW based on our analyses (Figs. 4a-d; Fig. S1-S6), although their population establishment scenarios were different. For example, our analyses suggested that Laos, Myanmar, Philippines, and PNG populations were likely linked to ‘bridgehead invasive founder China’s CY population’, the Vietnam population was more likely associated with individuals from Malaysia’s Johor State ‘bridgehead invasive founder population’ (Fig. 4C). DivMigrate analyses therefore suggested that populations in these SE Asian countries have diverse origins, both as potential direct recipients of New World FAW (e.g., Malaysia, also China) and as recipients of successfully established secondary invasive populations (e.g., Laos, Vietnam, Myanmar), with subsequent bi-directional movements between these countries.

Pest spread into Australia originating from Asia and SEA have been shown to be associated with wind patterns (Eagles et al. 2012), and separate introduction pathways for Western Australia and Queensland insect populations (e.g., the blue tongue virus vector *Culicoides brevitarsis*) have been reported (Tay et al. 2016). Similarly, arrival and subsequent detection of *S. frugiperda* populations across the northern region of Australia also likely represented separate establishments based on nuclear SNP analysis but did not support the reverse trajectory simulation findings of Qi et al. (2021). Populations between WA, NT, and Qld were shown to be genetically differentiated, suggesting limited gene flow between these populations at the early stage of invasions. The Kununurra population was found to be less genetically differentiated with the Johor (Malaysia) population (but also the population from India), suggesting a route moving south-east from South Asia/SE Asia into Western Australia, potentially via Indonesia. The Papua New Guinea population was shown to be not significantly differentiated from the Walkamin (Qld) population, while gene flow directionality analysis linked Yunnan populations (e.g., CY (Fig. 4a) and XP (Fig. 4b)) with the PNG population.

Boaventura et al. (2020) showed that Indonesian FAW individuals have unique insecticide resistance alleles for the AChE and VGSC genes. In China, Lv et al. (2021) reported diamide resistance alleles in individuals from Guangxi and Guizhou Provinces. While this study failed to source Indonesian and Cambodian populations (due to the difficulty of sharing and sending biological material from Indonesia and Cambodia to Australia), significant differences in resistance to indoxacarb was nevertheless detected between WA (Kununurra) and Qld (Strathmore) populations (Tay et al. 2021a, Tay et al. 2022c), and could be due to the different source populations for the WA and the Qld populations. On-going studies to monitor for potential arrivals of novel resistance alleles and those already reported in Indonesia and China to Australia will therefore be needed to inform and impact future FAW management strategies.

Phylogenetic analyses using both mitochondrial genomes and nuclear SNP loci supported multiple introductions at different scales. At least 12 C-strain mitochondrial genomes based on concatenated PCGs were identified, with some mitochondrial DNA genomes being detected only in SEA populations (e.g., Lao03 in Clade II, Lao12 in clade III). In the R-strain, at least 19 unique mitochondrial DNA genomes were detected (Fig. 1). Interestingly, and despite Australia representing the country with the most recent history of FAW’s incursions in this study, unique Australian mitochondrial DNA genomes not reported elsewhere in our other invasive populations were nevertheless detected (e.g., S3_Sf20-1 (Clade 1), KB1_107_AusRice (Clade III), 03_SF20-5_rice (Clade IV), 52_Sf20-1_Block2_21 (Clade V), and 55_Sf20-1_Block2_24 (Clade IX)). Some of these unique Australian mitogenomes shared close maternal evolutionary lineage with South Korean individuals (e.g., Clade III) that was further supported by nuclear SNP phylogeny (Fig. 2). Whether this represents linked anthropogenic-assisted invasion history remained to be confirmed, although direct flight between the two countries would be unlikely due to the large distance (*ca*. >7,000 km). Regardless, it highlights the challenge faced by biosecurity agencies globally in preventing accidental introductions of alien species.

An important finding from this study has been that the SE Asian populations were largely clustered away from other Asian (i.e., China, Indian) populations, supporting separate establishment histories of *S. frugiperda* populations in this region, and demonstrating the role played by independent, potentially anthropogenic-assisted introductions. The date of *S. frugiperda*’s establishment in China and SEA (e.g., Myanmar, Vietnam), though unknown, is potentially as early as pre-2014 (Gilligan and Passoa 2014; see also Vu 2008, Nguyen and Vu 2009, Tay et al. 2022d) and populations from SEA and China remained largely separate at least as recently as 2019/2020, based on analysis of our 870 SNP loci (see also Jiang et al. (2022) based on microsatellite DNA markers), contradicting the hypothesis of a Myanmar origin for the Yunnan *S. frugiperda* population (Sun et al. 2019, Wu Q et al. 2019, Li et al. 2020). Taken as a whole and incorporating population substructure (i.e., *F*_ST_) analysis, directional migration analysis of invasive *S. frugiperda* populations identified East Asia (e.g., China, South Korea) and SEA as biosecurity hotspots, while multiple introductions involving genetically distinct *S. frugiperda* likely occurred in north-western (i.e., Kununurra, WA) and north-eastern (i.e., Burdekin, Qld) Australia.

Population genomic studies via whole genome sequencing have enabled the spread patterns of this invasive pest to be re-evaluated to reveal highly complex scenarios that were likely associated with, or at least initiated by, trade activities. Whole genome sequencing is therefore proposed herein as essential for future biosecurity monitoring, in order to allow population genomics to directly impact on development of management policies and strategies (e.g., Guan et al. 2020), and to provide practical information to regulators and to some extent producers on how best to manage new invasive populations. The use of genome-wide SNP loci obtained via whole genome sequencing has enabled the elucidation of pathways to disentangle origins and establishment histories. This, bolsters risk assessment efforts, and provide information to biosecurity agencies and to international inter-governmental agencies to differentiate between natural and human-assisted spread. While it would be unlikely to stop natural migration of pests, accidental introductions via human-assisted spread are preventable. Findings from this study set the precedent that aims to inform biosecurity agencies and policy makers to effect behavioural change in the case where pest establishment and spread were underpinned by poor phytosanitary practices.

## Material and Methods

### FAW specimens for genomic analyses

*Spodoptera frugiperda* populations from the East Asia (i.e., South Korea), Southeast Asia (SEA; i.e., from: Myanmar, Laos, Vietnam, Malaysia, Philippines), and Pacific/Australia (i.e., Papua New Guinea, and Australia (representing new invasive range by *S. frugiperda*)) were surveyed for whole genome sequencing (Table S1; Fig. 6). Methods of specimen preservation involved collection of larvae/adults from fields and placing directly into highly concentrated ethanol (95-99.9%) and storing at -20°C until being used in DNA extraction. Samples sent to CSIRO from Queensland (CSIRO code: Sf20-1) and Western Australia (CSIRO code: SF20-4) were first used for bioassays purpose (Tay et al. 2021a, Tay et al. 2022c) prior to being stored at -20°C (once laboratory colonies were established) for DNA extraction purpose. Individuals from these two locations used therefore represent the original field-collected populations from northern Australia.

**Fig. 6:**
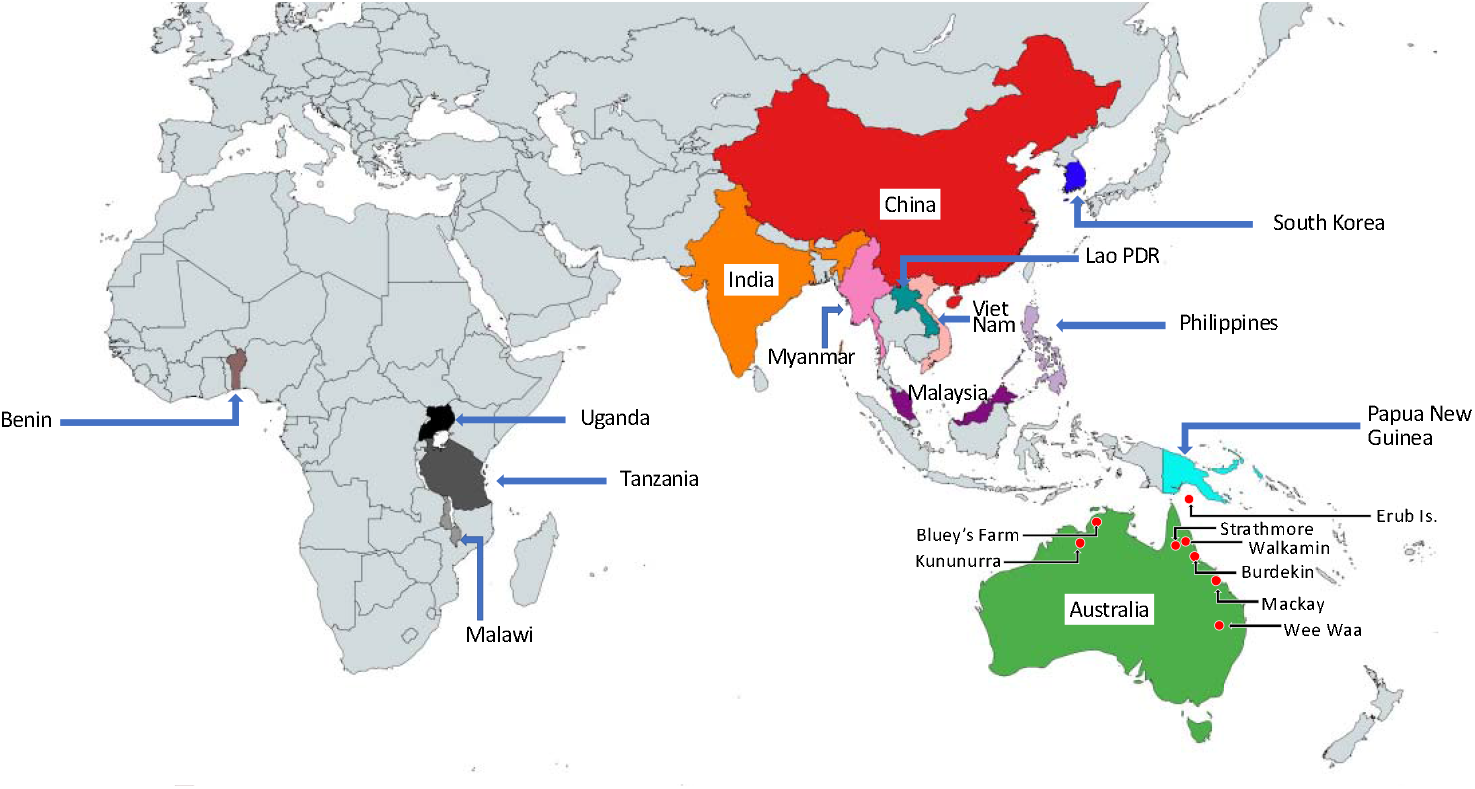
Countries where invasive populations of FAW surveyed for genomic analyses in this study were collected from (see main text, Table S1). Native New World populations (not shown) and invasive populations from Benin, Uganda, Tanzania, Malawi, India, and China have been reported in Tay et al. (2022d). Geographic regions of the sampled countries were as defined by World Atlas and are provided in Table S1.

### DNA extraction and genome library preparation

DNA of individual FAW samples (Table S1) was extracted using the Qiagen DNA extraction kit and eluted in 200μL elution buffer following the manufacturer’s protocol. Genomic DNA libraries for individual samples were prepared, quantified and sent to the Australian Genome Research Facility (AGRF) in Melbourne (Victoria, Australia) for commercial sequencing.

### Processing of genome sequences

Genome sequencing data for each individual were trimmed to remove adapter sequences using Trim Galore! (v 0.6.6; https://www.bioinformatics.babraham.ac.uk/projects/trim_galore/) and aligned to the *S. frugiperda* rice genome (v1.0) (Gouin et al. 2017) using bwa_mem2 (v2) (Vasimuddin et al. 2019). Duplicate alignments were removed using SAMBLASTER (v 0.1.26) (Faust and Hall 2014) and sorting completed using SAMtools (v1.9) (Li et al. 2009). Variants were then predicted using BBMap (v38.90) (Bushnell 2014) and normalised using bcftools (1.9) (Li et al. 2009). These variants were then subsampled using 870 SNP’s from Tay et al. (2022d) for further analyses.

### Evolutionary and population genomic analyses

For population and evolutionary genomic analyses of the SEA, South Korean, and Pacific/Australian FAW populations, we used the same set of the previously reported 870 neutral genome-wide SNPs (Tay et al. 2022d). We also included both native (i.e., Florida, Mississippi, Puerto Rico, Mexico, Guadeloupe, French Guiana, Peru, Brazil) and invasive (i.e., Benin, Malawi, Uganda, Tanzania, India, China) FAW populations that were reported in (Tay et al. 2022d) to infer the overall Maximum Likelihood (ML) phylogeny by IQ-Tree (Minh et al. 2013). Based on the phylogenetic analysis, we undertook separate admixture analyses (Alexander et al. 2009) by: (i) excluding New World, African, and Indian populations (i.e., include East Asia (China + South Korea) + Southeast Asia + Pacific/Australia), and (ii) excluding New World and Australian populations (i.e., including Africa + South Asia (India) + East Asia (Chinese, South Korea) + Southeast Asia + Papua New Guinea).

In the population genomic analysis, native FAW populations were excluded as the implication of multiple introductions that resulted in the establishment of Old World invasive FAW populations (e.g., Schlum et al. 2021, Tay et al. 2021b, Tay et al. 2022d) meant that inclusion of our limited native populations (i.e., from eight native localities) vs. the higher number of invasive FAW populations (i.e., from 13 invasive countries) were unlikely to provide greater certainty to detect potential origins of invasive populations. The study therefore uses 452 samples with an average sequencing depth of 19.24 (min. 8.71; max. 56.61; median 17.55) for evolutionary genomic interpretations relating to the gene flow patterns of the global FAW populations. Detailed description of the methods has been provided (Tay et al. 2021b, Tay et al. 2022d). Admixture was run using multiple K values (3 to 9), and the cross-validation error (CV) was quantified for each K. The K value with the least CV error (K=4 in this case) was selected for further interpretation. The 870 SNPs for this present study have been deposited in CSIRO public Data Access Portal (Rahul et al. 2022a).

### Population statistics

We used the populations program in Stacks (Catchen et al. 2013) to estimate the observed (Het_obs_) and expected (Het_exp_) heterozygosities for each population. The number of loci in the global population and individual populations departing significantly from Hardy-Weinberg equilibrium (HWE) was assessed using PLINK 2.0 (Chang et al. 2015). The populations program in Stacks was used to calculate nucleotide diversity (π) and *F*_IS_ (Wright’s (1951) inbreeding coefficient) and used only variant loci with no window size specified. We used Genepop (v4.7.5) (Raymond and Rousset 1995) to calculate pairwise comparisons of weighted *F*_ST_ values between populations, with significance of differentiation between populations tested using the exact G test.

### Mitochondrial Genome (mitogenome) analysis

The mitogenomes of all individuals from SEA (n=94), East Asia (South Korea, n=1<u>2</u>), and Pacific/Australia (n=164) were individually assembled following the approach as described in Tay et al. (2022d). Briefly, we used Geneious 11.1.5 to map raw sequences for each FAW individual against a reference genome (GenBank accession number MT897262 (for mtCOI C-strain)/MT897275 (for mtCOI R-strain); Tay et al. 2022d), followed by fine-tuning to resolve regions of poor assembly and ambiguity. Fully assembled mitogenomes were then annotated using MITOS (Bernt et al. 2013) by selecting invertebrate mitochondrial genetic code 5. Bed files from MITOS annotations were imported into Geneious 11.1.5 to fine-tune the predicted annotations of gene and protein coding genes (PCGs).

All FAW mitogenomes from this study and those from Tay et al. (2022d) were combined and aligned using MAFFT Alignment v7.450 (Katoh et al. 2002, Katoh and Standley 2013) within Geneious 11.1.5, using the default Auto option for Algorithm, 1.53 for Gap open penalty 0.123 for offset value, and 200PAM / K = 2 for Scoring matrix. Once aligned, we trimmed and removed all intergenic regions including the A-T rich replication origin, all tRNAs, and the two rRNA genes, leaving 11,303 bp of the 13 concatenated PCGs. We identified unique concatenated PCG mitogenomes within individual countries using the DNAcollapser in FaBox (1.5) (Villesen 2007), and partitioned these unique concatenated PCG mitogenomes prior to inferring ML phylogenies without outgroups using IQ-tree implementing UFBoot (Minh et al. 2013) with 1,000 replications to ascertain node confidence. Visualisation of the phylogeny was by Dendrogram (Huson and Scornavacca 2012) to enable a conservative estimate of the number of maternal lineages in the invasive populations. All assembled and annotated mitogenomes used have been deposited in CSIRO public Data Access Portal (Rahul et al. 2022b).

### Strain assessment

With the non-recombinant nature of the mitogenomes, we analysed the mitogenomes of the FAW according to their strain classification (i.e., C-strain (corn strain) and the R-strain (rice strain)) separately to identify the conservative number of maternal founders for both strains across the invasive ranges, based on concatenation of the 13 PCG regions only, excluding the tRNA and rRNA genes, and the variable intergenic and AT-rich regions. We follow the naming suggestion of Tay et al. (2022b) and refer the rice-strain and corn-strain instead as R-strain and C-strain, respectively. Individual FAWs were characterised as the R- or C-strain by comparing their relevant partial mt*COI* gene regions against previously characterised reference sequences (rice (R-strain): GenBank MF197867; corn (C-strain): GenBank MF197868; see Otim et al. 2018b). We note that classification of the FAW host-strain (i.e., R-strain/C-strain) is contentious depending on the markers used (e.g., Dumas et al. 2015 vs. Nagoshi 2010) and may have limited applications in hybrid populations (Tay et al. 2022b, Tay et al. 2022d), we nevertheless provided this information based on both the partial mt*COI* gene identity and the *TPI* exon 5 gene region as previously described (i.e., for the *TPI* loci, to differentiate between the C-strain and R-strain *S. frugiperda*, whole genome sequence data were mapped to the SFRU_RICE_002481 contig (Gouin et al. 2017) at location 12939.

### Principle Component Analysis (PCA)

Principal Component Analysis (PCA) was also carried out to increase interpretability of the large and complex FAW genomic dataset through maximising variance by creating new uncorrelated variables and to aid in visualisation. PCA based on the 870 SNPs was performed using PLINK v1.9 (Purcell et al. 2007) and visualised using ggplot2 (Wickham 2016). To enable visualisation of evidence of independent introductions we compared (i) China (CY, YJ, XP) populations vs. Myanmar, (ii) within Malaysian populations between those collected from Penang and Johor States vs. Kedah State; (iii) between China and East Africa (e.g., Uganda, Malawi), and (iv) between China, Malaysia (Kedah State), and Australia (NT, NSW)).

### DivMigrate analysis

DivMigrate analysis was carried out using the complete set of 870 SNP loci for all invasive populations from Africa, Asia, and SEA, with the exception of the Kedah State (Malaysia) population that showed unique genomic background (and represented a population from the on-going lab-maintained culture first collected from maize field in 2019). Australia populations representing the ‘final arrival’ populations, were also excluded in the DivMigrate analysis to increase interpretability relating to East Africa, South Asia, East Asia and SEA populations. Relative migration was assessed using the G_ST_ statistics with 100 bootstrap replications and filter threshold (i.e., visualisation of the number of links in the network plot) was set at 0.453 and plotted using the graph visualisation and manipulation software Gephi 0.9.2 <www.gephi.org> and countries were separately drawn using Microsoft PowerPoint for Mac v16.54.

## Acknowledgements

We would like to thank Dr Brian Thistleton (Department of Industry, Tourism and Trade, NT), Dr Ian Newton and Dr Melina Miles (QDAF), Mr Jack Daniel (Northern Australia Crop Research Alliance Pty Ltd, Kununurra WA), Prof. Myron Zalucki (University of Queensland), Mrs Anna Madden (Wee Waa, NSW), and Dr Simone Heimoana (CSIRO), for providing Australian FAW specimens for genomic analyses. The Erub Island FAW was kindly provided by the Department of Agriculture, Water and the Environment (DAWE) and Northern Australia Quarantine Strategy (NAQS). Van NTH and Huyen NPT (PPRI, Vietnam) helped with FAW specimen and related metadata collection. DR. Donald Kachigamba provided samples from Malawi. Bill James (CSIRO) helped with laboratory work, Dr Sharon Downes, Bill James, and Dr Sarina Macfadyen (CSIRO), and Dr Eric Huttner (ACIAR) provided helpful discussion during the course of this work. The work performed at INRAE was publicly funded through the ANR (the French National Research Agency, Grant ID 1702-018, given to K.N.) under the “Investissements d’avenir” programme with the reference ANR-10-LABX-001-01 Labex Agro and coordinated by Agropolis Fondation under the frame of I-SITE MUSE (ANR-16-IDEX-0006). It was also funded by a grant from the department of Santé des Plantes et Environnement at Institut national de la recherche agronomique for K.N. (adaptivesv), and was also financially supported by EUPHRESCO (FAW-spedcom, given to Anne-Nathalie Volkoff). The project led by CSIRO is a co-investment by the Grains Research and Development Corporation (GRDC), the Australian Centre for International Agricultural Research (ACIAR), the Cotton Research and Development Corporation, FMC Australasia and Corteva Agriscience. GRDC manages the project on behalf of the co-investors.

## Data availability

All data used in this study have been deposited in the CSIRO Data Access Portal and are freely accessible, under the permanent links https://doi.org/10.25919/24gy-g132 and https://doi.org/10.25919/c53m-ts57.

