## Supplementary material for "Complex multiple introductions drive fall armyworm invasions into Asia and Australia": Table S1

**Table S1:** Population data of invasive *Spodoptera frugiperda* samples from Pacific/Australia (PA; Papua New Guinea, Australia), Southeast Asia (SEA; Vietnam, Lao PDR, Philippines, Myanmar) and South Korea (East Asia) used in whole genome sequencing for population genomic analysis. Number of specimens from each location is indicated by ‘N’. Where available, GPS co-ordinates and elevation are provided. The number of individuals belonging to R- or C-strain mitochondrial DNA *COI* (mt*COI*) haplotypes is also provided. Annotated mitochondrial DNA genomes from this study in GenBank format (.gb) is can be downloaded from CSIRO public Data Access Portal (Rane et al. 2022b).

| **Country** | **State/Province** | **Population** | **N** | **Date Collected** | **note** | **R: C** |
| --- | --- | --- | --- | --- | --- | --- |
| Australia | Qld | Walkamin Research Station | 26 | 6-24 Apr-20 | Details accompanied samples as: ‘Block 1 unsprayed’ (n = 8), ‘Block 1 sprayed 24Apr’ (n = 10), ‘Block 2 Early sprayed’(n = 9), ‘Block 3, near Mango’ (n = 2) | 26 : 0 |
|  | Qld | Burdekin | 29 | 11-Mar-20 | ‘Sf20-2’ Home Hill, Maize host | 29 : 0 |
|  | Qld | Strathmore | 30 | 03-Mar-20 | ‘Sf20-1’; see Bioassay report for details | 30 : 0 |
|  | Qld | Mackay | 7 | 29-Jul-20 | Corn field, Marian Qld. | 7 : 0 |
|  | WA | Kununurra | 15 | 17-29 May-20 | Samples ‘KB1’ (n = 8) GPS: -15.781658, 128.719078; ‘KB2’ (n = 7) GPS: -15.781658, 128.719078 | 15 : 0 |
|  | WA | Kununurra | 17 | 16-Aug-20 | 'Sf20-4', G_0_; 9 other larvae as G_8_ (May-2021) | 17 : 0 |
|  | NT | Bluey's Farm | 6 | Jun-20 | ‘Sf20-5’ | 6 : 0 |
|  | NSW | Wee Waa | 8 | Nov-20 | Parasitised by *Cotesia ruficrus* | 8 : 0 |
|  | QLD | Erub Island | 1 | 20-Jan-20 | Pheromone traps | 1: 0 |
| South Korea | Milyang (MF) | South Korea (N = 12) | 7 | Feb-20 | Milyang-si, Gyeongsangnam-do, G_4_ lab, GPS: 35˚29.29 N, 128˚44.31 E | 7 : 0 |
|  | Goryong (GR) |  | 1 | Sep-19 | Goryeong-gun, Gyeongsangbuk-do, Corn-field; GPS: 35˚ 64.04 N, 128˚39.08 E | 1 : 0 |
|  | Haenam (MN) |  | 1 | Aug-19 | Haenam-gun, Jeollanam-do, Corn field; GPS: 34˚24.38 N, 126˚37.55 E | 1 : 0 |
|  | Milyang (MY) |  | 2 | Sep-19 | Milyang-si, Gyeongsangnam-do, Corn field; GPS: 35˚29.29 N, 128˚44.31 E | 0 : 2 |
|  | Muan (MA) |  | 1 | Aug-19 | Muan-gun, Jeollanam-do, Corn field; GPS: 34˚52.21 N, 126˚31.11 E | 1 : 0 |
| Vietnam |  | Vietnam | 11 |  |  | 6 : 5 |
|  | Nghe An | Nam Dan |  | 5-March 2020 | Nam Dan, Nghe An, corn field, GPS: 18.69527 N’, 103.50277 E |  |
|  | Nam Dinh | Nam Truc |  | 28 April-2020 | Nam Truc, Nam Dinh, Corn field, GPS: 20.33027 N, 105.59083 E |  |
|  | Ha Noi | Gia Lam |  | 16-June-2020 | Gia Lam, Ha Noi, corn field, GPS: 20.93916 N, 105.89083 E |  |
|  | Lao Cai | Sa Pa |  | 11-June-2020 | Sa Pa, Lao Cai, corn field; GPS: 22.34611 N, 103.84222 E |  |
|  | Hoa Binh | Cao Phong |  | 17-June-2020 | Cao Phong, Hoa binh, corn field, GPS: 20.70138 N, 105.33000 E |  |
|  | Ha Noi | Soc Son |  | 19-June-2020 | Soc Son, Ha Noi, corn field; GPS: 21.28527 N, 105.76666 E |  |
|  | Hoa Binh | Hoa Binh town |  | 24 June-2020 | Hoa Binh town, Hoa Binh, corn field; GPS: 20.82277 N, 105.33361 E |  |
|  | Ha Noi | Dan Phuong |  | 22-July- 2020 | Dan Phuong, Ha Noi, Corn field, GPS: 21.11472 N, 105.61638 E |  |
|  | Ninh Binh | Tam Diep |  | 26-July-2020 | Tam Diep, Ninh Binh; corn field; GPS: 20.7841 N, 105.82972 E |  |
|  | Son La | Van Ho |  | 25-July-2020 | Van Ho, Son la, corn field; GPS: 11.79194 N, 104.76000 E |  |
|  | Ha Noi | Dong Anh |  | 21-August-2020 | Dong Anh, Ha Noi, GPS: 20.7841 N, 105.82972 E |  |
| Lao PDR | Vientiane | Vientiane | 4 | 23-Feb-20 | Koksay village, Don village, Nahai village, Hatxayfong district, Vientiane capital, GPS: 17′.8137, 102′.6866; GPS: 17′.8378, 102′.6132; GPS: 17′.87.19.51, 102′.16.34.69; GPS: 17′.5954967 ,102′.5922925 | 1 : 3 |
|  | Xiengkhouang | Xiengkhouang | 10 | 7-Jul-20 | Khangphanien village, Yordhuay village, Nammen village ; Nonghat district, Xiengkhouang province, GPS: 19′.5886, 103′.78354; GPS: 19′.59894, 103′.78365; GPS: 19′.58610, 103′.73989  Napan village, Khaivieng village, Phousun village Kham district, Xiengkhouang province, GPS: 19′.61025, 103′.58940; GPS: 19′.61020, 103′.58942; GPS: 19′.64898, 103′.51767 | 7 : 3 |
|  | Champasak | Champasak | 4 | 11-14 Jul-20 | Thongsadam village, Khong district, Champasak province, GPS: 19′.60027, 103′.43980  Huayko village, Nakham village, Phaolamphan village Pathoumphone district, Champasak province, GPS: 14′.79028, 106′.01612; GPS:14′.86192, 105′.93782; sampleFW14 GPS:14′.86192, 105′.93782  Don village, Kaengkia village, Phonphai village Bachieng district, Champasak province, GPS: 15′.267111, 105′.93223; GPS: 15′.40315, 106′.08103; sample FW19 GPS: 15′.22560, 106′.01612  Kaper village, Sekapoung village Paksong district, Champasak province, GPS: 15′.28314, 106′.19620; GPS: 15′.25843, 106′.23049 | 4 : 0 |
| Philippines | Batangas | Lipa City | 20 | 3, 5, 20-July-20 | Corn field; 13° 56' 30.754" N 121° 9' 51.912" E | 14 : 6 |
| Malaysia | Johor | Labis | 10 | 22-Sep-20 | GPS coordinates not provided | 7: 3 |
|  | Penang | Seberang Perai | 9 | 8-Oct-20 | GPS coordinates not provided | 3 : 6 |
|  | Kedah | Changlun | 9 | Sep-19 | Laboratory population established in 2019 from field-collected samples from Changlun, Kedah | 9 : 0 |
| Myanmar | Southern Shan State | Aungban Research Farm (AB) | 3 | Sep-20 | 20˚40'40" N 96˚40'39" E ; Altitude: 1259m | 2 : 1 |
|  | Magwe Region | Kimpoungtaung Research Farm (KPT) | 3 | Sep-20 | 20˚01'53" N, 95˚22'22" E ; Altitude: 102m | 2 : 1 |
|  | Eastern Shan State | Kengtung Research Farm (KT) | 3 | Sep-20 | 21˚17'25" N 99˚38'53" E ; Altitude: 796m | 1 : 2 |
|  | Naypyitaw | Tatkone Research Farm (TK) | 2 | Sep-20 | 20˚08'21" N 96˚12'43" E ; Altitude: 145m | 1 : 1 |
|  | Naypyitaw | (WAX) | 3 | Sep-20 | 19˚49'06" N 96˚16'00" E ; Altitude: 96m | 2 : 1 |
|  | Naypyitaw | Yezin (YZ) | 3 | Sep-20 | 19˚49'32" N 96˚16'48" E ; Altitude: 98m | 1 : 2 |
| Papua New Guinea | Madang Province | Ramu Sugar Estate | 16 | 15-17 Jun-2020 | 5°58.154 S, 145°53.252 E, Maize host | 16 : 0 |
|  | Central Province | Yule Island Junction | 1 | 5-Jun-20 | corn host | 1 : 0 |

**Note:** We followed World Atlas definition for sub-Saharan Africa, East Asia (China, Japan, Koreas, Taiwan), South Asia (Afghanistan, Pakistan, India, Bangladesh), Southeast Asia (Brunei, Cambodia, Indonesia, Laos PDR, Malaysia, Myanmar, the Philippines, Thailand, Timor-Leste, Vietnam) and Australia/Pacific (Papua New Guinea, Australia, New Zealand, Pacific Islands).
