## Supplementary material for "Complex multiple introductions drive fall armyworm invasions into Asia and Australia": Figure S1-S6

**Supplemental Figures**

**Figure S1-S6:** DivMigrate analysis of invasive *Spodoptera frugiperda* populations from Africa (Benin (BEN), Uganda (UGA), Malawi (MWI)), South Asia (India (IND), East Asia (China (CHN), South Korea (KOR)), Southeast Asia (Myanmar (MMR), Laos People Democratic Republic (LAO), Viet Nam (VNM), Philippines (PHL), Malaysia (MYS)), and Papua New Guinea (PNG). Multiple populations from the Yunnan province in China (Cangyuang (CY), Yuanjiang (YJ), Xinping (XP)) and from the Penang (PN) and Johore (JB) states in Malaysia were included. Estimates of migration rates were calculated using G_ST_ with significant rates (shown as red values) at alpha = 0.5 estimated from 100 bootstrap replications. Note that no migration events were established for Benin and South Korea, suggesting unique introduction events among the invasive populations included in this analysis. Australian and Malaysia Kedah state populations were excluded to enable ease of interpreting migration patterns. See main text for detailed discussion.

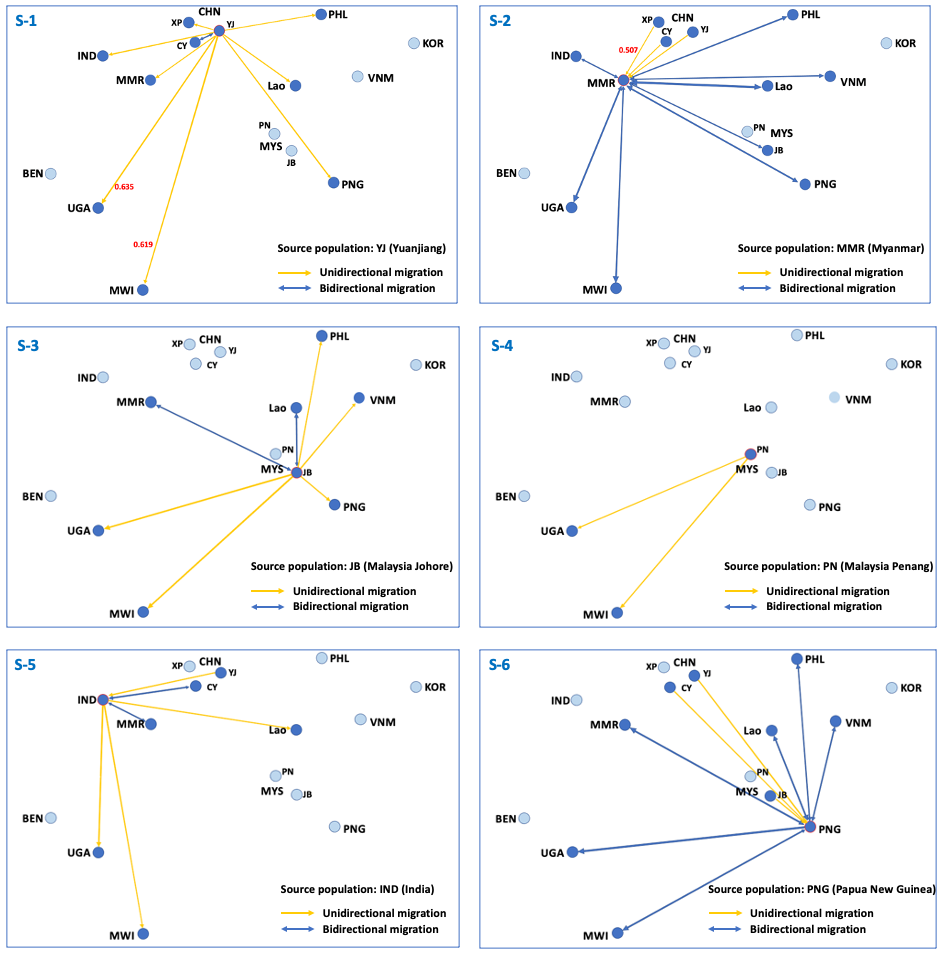
